# Confidence as a priority signal

**DOI:** 10.1101/480350

**Authors:** David Aguilar-Lleyda, Maxime Lemarchand, Vincent de Gardelle

## Abstract

When dealing with multiple tasks, we often find ourselves in the problem of establishing the order in which to tackle them. Here we asked whether confidence, the subjective feeling in the accuracy of our decisions, plays an active role in this ordering problem. In a series of experiments, we show that confidence acts as a priority signal when ordering responses about tasks already completed, or ordering tasks that are to be made. In experiments 1-3, participants were engaged in a dual task and categorized perceptual stimuli along two dimensions. We found that they tended to give first the decision in which they were more confident. We also prove that confidence drives prioritization above and beyond task difficulty or response accuracy, and we discard alternative interpretations in terms of response availability or task demands. In experiments 4-6, we show that when participants have to select which of two sets of trials they want to perform first, they engage first in the set associated with higher confidence, and we extend this finding to situations involving non-perceptual (mental calculation) decisions. Our results thus support the role of confidence as a priority signal, thereby demonstrating a new way in which it regulates human behavior.

**Highlights:**

1. We show that when having to decide the order in which to approach two tasks, humans prefer to start with the one they feel more confident in.
2. This holds both when deciding in which order to report two already completed tasks, and when deciding the order in which to tackle two tasks yet to complete. Our results are replicated in perceptual and non-perceptual situations.
3. The role of confidence on prioritization cannot be reduced to that of task difficulty or response accuracy.
4. Our findings demonstrate a new way in which confidence regulates human behavior.

---

It is frequent to start a day at work by compiling in a to-do list all the tasks to complete that day. But once the list is written, where do we start? Which task shall we tackle first, and which tasks can be postponed? Of course, this problem is sometimes solved by taking into account external constraints, such as scheduled meetings, imminent deadlines, or the limited availability of specific tools or collaborators. In other circumstances, however, we are free to decide in which order to complete our tasks. In such unconstrained situations, humans may not perform their tasks in a random order, and they might instead exhibit some systematic preference for doing one task before another. Here, we suggest that confidence may play a role in this prioritization problem. In a situation where two tasks have to be done, individuals may address first the one about which they feel more confident. In a situation where two completed tasks have to be reported, individuals may start by the one in which they were more confident. In both situations, confidence would be a priority signal.

Different considerations make this hypothesis plausible. Firstly, even in the absence of external rewards, the feeling that we have completed a task successfully, and that we can cross an item off our to-do list, may be intrinsically rewarding. Confidence corresponds to this feeling of success in the task. This feeling is generally valid in perceptual^1,2^ and memory tasks^3^, where confidence typically correlates with performance (although some dissociations have been documented^4–6^). Therefore, in order to secure rewards as soon as possible, people could seek to complete first the task in which they are more confident^7^. Another reason to expect that humans would use confidence as a priority signal is to reduce their mental workload^8^ before facing the most demanding tasks. Indeed, having multiple pending tasks can diminish our cognitive resources, so to avoid facing the most demanding tasks with a reduced mental workload, agents could complete first the less demanding tasks, in which they are more confident. To do so, they could compare their confidence across the different tasks at hand^9,10^, and prioritize them accordingly.

We evaluated whether confidence serves as a priority signal in a series of experiments involving elementary decisions. Our first experiment aimed at evaluating whether confidence would relate to the prioritization of responses between two perceptual decisions already made. On each trial (see Figure 1A), participants (n=28) were presented an array of colored letters and they had to report both the dominant color (orange or blue) and the dominant letter (X or O) in the stimulus. Immediately after they had to give a confidence rating for each choice, on a subjective probability scale of success ranging from 50% (i.e. pure guess) to 100% (i.e. completely certain of being correct). By calibrating the stimulus before the main experiment (see methods), we obtained for each task an easy and a hard condition, which corresponded to an expected 90% and 60% of correct responses, respectively. As Figure 1B shows, observed performance in the experiment closely matched expected performance.

**Figure 1.**
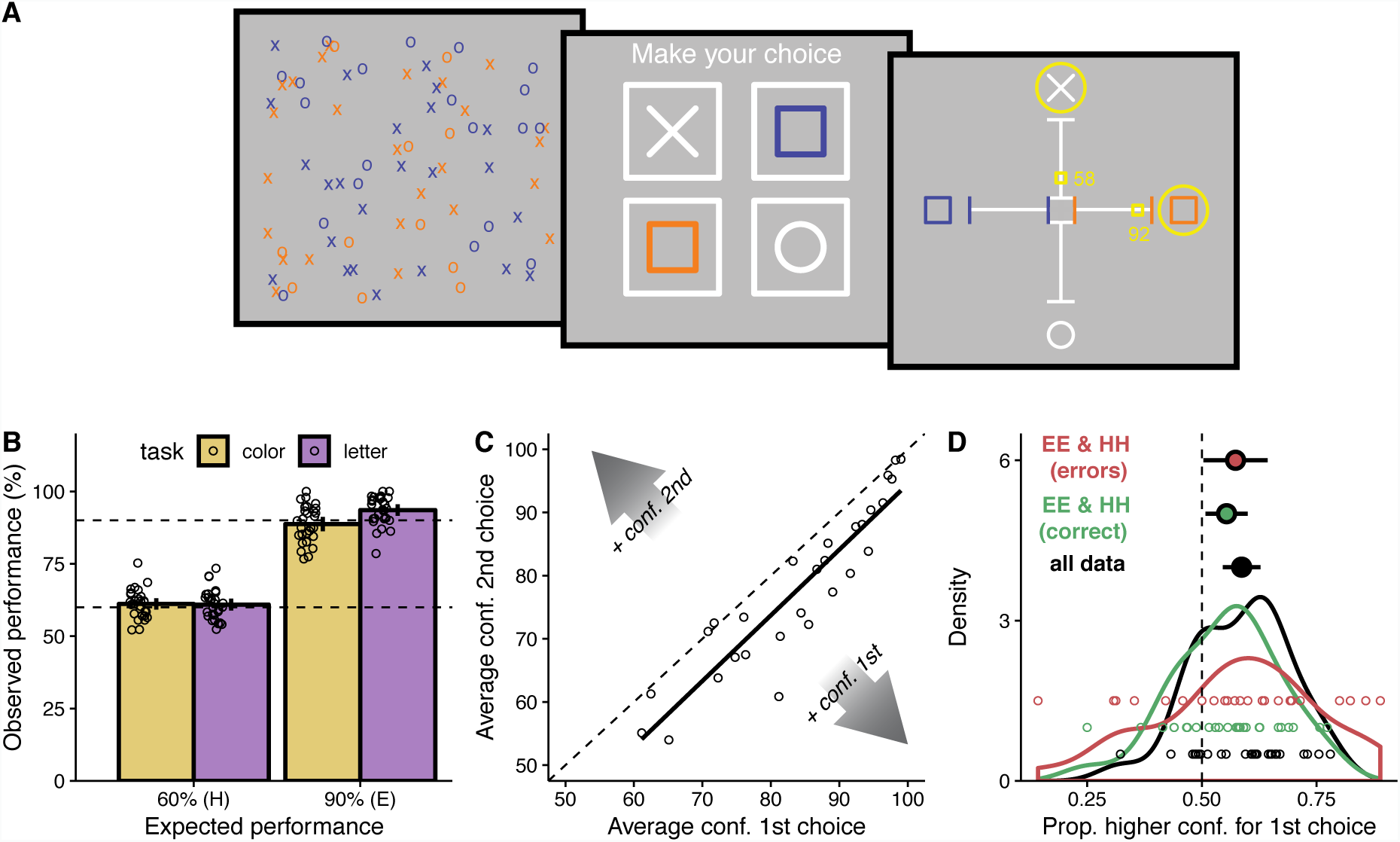
Experiment 1: design and results. **A.** Overview of a trial. An array of 80 elements appeared for 1s. Each element could be an O or an X, colored blue or orange. After the stimuli disappeared, a response screen appeared. Four boxes were presented, each containing one of the four elements related to the choice. Participants gave their response to the color task by clicking on the box containing either the blue or the orange square, and to the letter task by clicking on the box containing either the O or the X. Next two scales, one for the color task and another for the letter task, appeared on the screen disposed as a cross. Participants could then rate their confidence on each task’s decision, from 50 (total uncertainty) to 100 (total certainty). **B.** Observed performance as a function of expected performance, both in percentage, split by task type. Bars represent averages across participants, with error bars denoting 95% confidence intervals. Dots represent individual participants. Top and bottom dashed lines help indicate where 90% and 60% observed performance would lay, respectively. **C.** Average confidence rating given for the second choice within a trial, as a function of that for the first choice. Dots represent individual participants, and the black, solid line represents a linear fit of the data. Participants below the dashed, identity line expressed more confidence in their first choices, and vice versa. **D.** Density of the proportion of trials where the first choice had a higher confidence rating than the second choice. The black density corresponds to the whole dataset, the green density corresponds only to those EE and HH trials where both choices were correct, and the red density corresponds only to those EE and HH trials where both choices were errors. Densities are drawn by using each participant’s data, with each participant’s average shown as an individual dot at the bottom. Across-participant averages are displayed as big dots with error bars showing 95% confidence intervals. The dashed line corresponds to equal confidence ratings for first and second choices.

Critically, although participants were not asked to report their judgments in a specific order, we expected that they would report first the judgment associated with a higher confidence. This was indeed the case on average across trials: confidence in the first choice was higher than confidence in the second choice (Figure 1C, t(27) = 6.425, *p* < 0.001, 95% CI [4.166, 8.075], *d* = 1.214). It was also the case within each trial: as seen in Figure 1D, the proportion of trials in which the confidence associated to the first choice was greater than that associated to the second was systematically greater than 0.5 (t(27) = 4.339, *p* < 0.001, 95% CI [0.546, 0.628], *d* = 0.820). Furthermore, this pattern held even when the two perceptual dimensions had the same difficulty level and responses were both correct (green line in Figure 1D, t(27) = 2.400, *p* = 0.024, 95% CI [0.508, 0.600], *d* = 0.454) or both incorrect (red line in Figure 1D, t(27) = 2.150, *p* = 0.041, 95% CI [0.503, 0.643], *d* = 0.406). Note that, by construction, this later analysis confirms that participants’ priority was driven by confidence per se, and not simply by task difficulty or response accuracy.

To strengthen this point, we conducted four more analyses on this data, which showed that confidence affects priority above and beyond task difficulty (expected accuracy) and actual accuracy. First, in a logistic regression model, we found that responding to the color task first was significantly predicted from the difference in reported confidence between the two tasks (*β* = 0.045, S.E. = 0.002, *p* < 0.001), even when the difference in actual accuracy (*β* = 0.002, S.E. < 0.001, *p* < 0.001) and in expected performance (*β* = 0.003, S.E. = 0.001, *p* = 0.004) between the two tasks were included as predictors in the same regression. Second, we compared this regression model with a simpler one that only included the difference in expected performance and actual accuracy between tasks, and we found that including confidence significantly improved the model (log-likelihood: full model = −6083.624, no confidence model = −6644.367; χ²(1) = 1121.486, *p* < 0.001). Third, in a different but similar approach, we could predict priority from confidence (*p* < 0.001) in a regression in which we included as an offset the prediction of priority by expected performance and actual accuracy. These analyses were conducted on the full data and are presented in more details in supplementary information 4. In our last analysis our goal was to pit confidence against accuracy by evaluating whether participants would prioritize an error made with greater confidence over a less confident but correct response. To do so, we focused on a narrow subset of trials where participants made an error for one dimension and a correct response for the other, but expressed a greater confidence in their error than in their correct response. Pooling all participants together (due to the limited number of trials per participant), we performed a binomial exact test and found indeed that the proportion of trials where the error was prioritized was greater than 0.5 (actual proportion = 0.570, n = 1384 trials, *p* < 0.001). Results were similar if only taking trials where the two dimensions had the same difficulty level (proportion = 0.591, n = 658 trials, *p* < 0.001). Thus, when confidence and accuracy were dissociated, the former still drove prioritization of responses. This not only reinforces the idea that the role of confidence cannot be reduced to that of accuracy, but also suggests that the role of confidence is preponderant.

In the experiment described above, participants may have associated their confidence ratings with task priority simply because they had to explicitly report their confidence. Our result (i.e. ‘most confident response first’) would be strengthened if replicated with a paradigm that eliminated this possibility of an implicit demand effect. In our second experiment, thus, we tested our hypothesis without asking for confidence ratings, and we relied instead on task difficulty as a proxy for confidence. Indeed, in our first experiment we observed that reported confidence was greater for easy tasks than for hard tasks, and also greater for correct responses than for errors (for details see supplementary information 3.2.), as typically found in the confidence literature^1,3,11^.

In Experiment 2, we therefore asked a new set of participants (n=20) to perform the same dual task without confidence ratings. Figure 2A depicts the proportion of times the first response was related to the color task, in each of the four possible kind of trials: easy color - easy letter (EE), easy color - hard letter (EH), hard color - easy letter (HE), and hard color - hard letter (HH). If confidence truly guided the priority of the responses, participants should exhibit a bias towards responding to the easy task before the difficult task. We found that, indeed, the color judgment was reported first more in EH trials than in HE trials (42% vs. 37% of trials). This difference in proportion was highly consistent across participants (t(19) = 3.059, *p* = 0.006, 95% CI [0.018, 0.097], *d* = 0.684), as illustrated by Figure 2B. In other words, participants’ task prioritization was affected by task difficulty, as a proxy for confidence, such that participants exhibited an easy-first bias. For completeness, we also confirmed that the same results were obtained in Experiment 1 (see supplementary information 3.3.).

**Figure 2.**
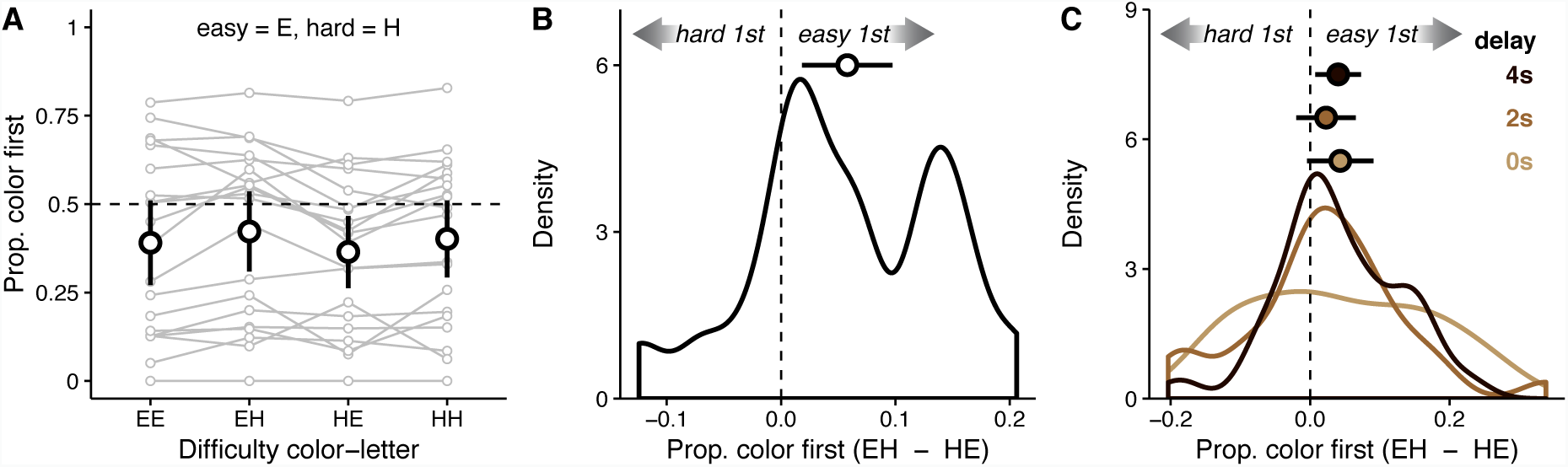
Experiment 2 & Experiment 3: results. **A.** For Experiment 2, proportion of trials where the color task was first responded to, as a function of the type of trial (EE = easy color, easy letter; EH = easy color, hard letter; HE = hard color, easy letter; HH = hard color, hard letter). Each set of four dots connected by lines represents one participant. Bigger dots represent average proportions across participants, and error bars denote 95% confidence intervals. The dashed line corresponds to no preference towards responding first to the color or the letter task. **B.** For Experiment 2, density of the difference between proportion color task chosen first for EH trials and proportion color task chosen first for HE trials. The density is drawn by using the difference of proportions for each participant. The across-participant average is displayed as a big dot with error bars showing 95% confidence intervals. The dashed line corresponds to no difference between EH and HE trials. **C.** For Experiment 3, density of the difference between proportion color task chosen first for EH trials and proportion color task chosen first for HE trials. The three densities, one for each delay, are drawn by using the difference of proportions for each participant. Across-participant averages are displayed as big dots with error bars showing 95% confidence intervals. The dashed line corresponds to no difference between EH and HE trials.

In our third experiment, we wanted to discard one possible interpretation of our results in terms of response availability. Because in our previous experiments the response screen was presented immediately after the stimuli disappeared, one could argue that only the decision for the easy task had been reached by then. If that was true, then the easy-first priority effect could be due to response availability rather than confidence. To tackle this concern, we modified our previous experiment by introducing a delay (0s, 2s, or 4s, randomized across trials) between the stimulus offset and the onset of the response screen. We reasoned that, if the easy-first effect was only driven by response availability at the response screen onset, then it should disappear for long delays. We ran this experiment with new participants (n=30) and then calculated again the proportion of trials in which color was first responded to in EH trials and in HE trials, now for each delay condition. We found that the main effect of trial type was significant overall (*F*(1,29) = 6.098, *p* = 0.020, 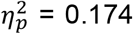), with an easy-first bias replicating our previous studies. Although the main effect of delay was also significant (*F*(2,29) = 12.760, *p* = 0.020, 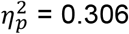), it did not interact with trial type (*p* = 0.679, see also Figure 2C). This data thus ruled out the concern that participants responded to the easy task first because it was the only task for which a decision was available by the time the response screen was presented.

The experiments described so far showed that confidence acts as a priority signal at the level of response execution: when engaged in a dual task, participants tend to communicate first the response associated with greater confidence. In what follows, we investigated the prioritization between two tasks, instead of between responses. In other words, in a situation where two tasks have to be completed separately (much like our introductory “to-do list” example), we asked whether participants would engage first in the task that they feel more confident about. In such situations, participants may rely on prospective confidence, that is their anticipation about the probability of success in a task before they actually do it, which can be based for instance on prior experience with the same task^12,13^. We implemented this idea in our next experiments. First we familiarized participants with two conditions, one eliciting higher confidence than the other. Then in a subsequent test phase, they would face again the same two conditions, in the order they preferred, and we evaluated whether they would prioritize the condition associated with higher confidence.

Our fourth experiment was conducted on a new sample of participants (n=31), which completed 32 experimental blocks, each constituted of a familiarization and a test phase (Figure 3A). In the familiarization phase, participants completed two sets of trials. Each set contained 6 trials, and was associated with a name and a task (either color or letter). Importantly, one set contained only hard trials, while the other set contained only easy trials. Difficulty was calibrated before the experiment (Figure 3B), and the order of the easy and hard set in the familiarization phase was random. After completing the familiarization phase, the test phase started: participants were presented with the names of both sets, and they could choose the order in which they wanted to complete 4 trials of each set. They knew that performance in these test trials would determine their final payoff. Critically, participants chose more often to face the easy set first (Figure 3C, t(30) = 2.284, *p* = 0.030, 95% CI [0.503, 0.557], *d* = 0.410). These results thus demonstrate that prospective confidence too can establish the priority of one task over another. Interestingly, whether the two sets involved different tasks or the same task did not affect this priority effect (*p* = 0.256). This suggests that the comparison of the two sets involved relatively abstract representations, as was found for confidence comparison^9,14^.

**Figure 3.**
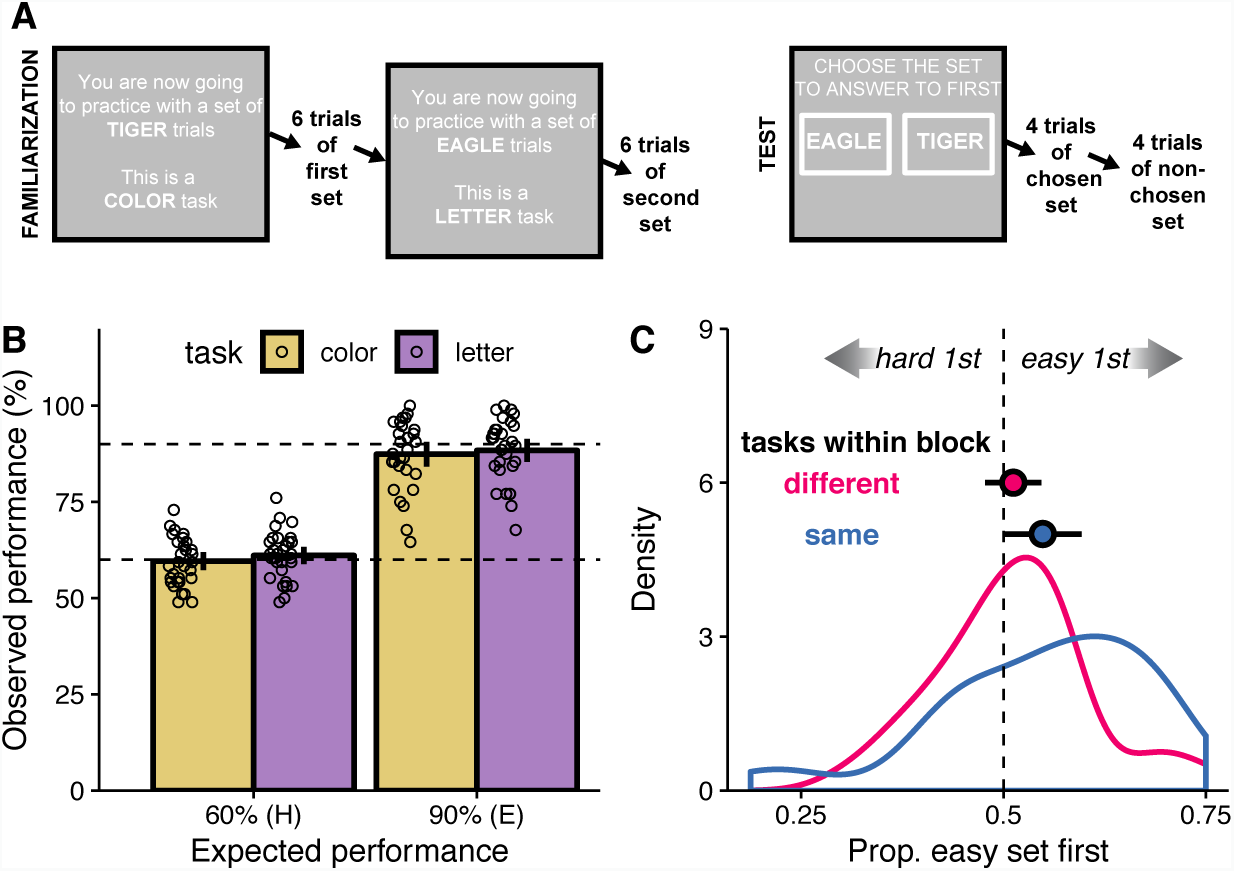
Experiment 4: design and results. **A.** Overview of one block. The familiarization phase made participants practice with 6 trials of two different sets, with each set introduced by a screen stating its name and the type of task participants would have to respond to. During the test phase participants had to complete 4 trials of each set. At the beginning of the phase, a screen with the names of both sets was presented, and participants clicked on the name of the set they wanted to start with. **B.** For the familiarization phase, observed performance as a function of expected performance, both in percentage, split by task type. Bars represent averages across participants, with error bars denoting 95% confidence intervals. Dots represent individual participants. Top and bottom dashed lines help indicate where 90% and 60% observed performance would lay, respectively. **C.** Density of the proportion of blocks where participants chose to complete first the 4 trials from the set with an easy task. One of the displayed densities corresponds to blocks where both tasks were the same, the other to those where they were different. Both densities are drawn by using each participant’s data. Across-participant averages are displayed as big dots with error bars showing 95% confidence intervals. The dashed line corresponds to easy and hard tasks being chosen first equally often.

In Experiment 5 (n=29), our goal was to further demonstrate that task prioritization between two sets of trials depends on confidence above and beyond performance. To do so, we reproduced the structure of Experiment 4 while dissociating confidence from performance, by capitalizing on previous work showing that confidence can be biased by the variability of the evidence available for the perceptual decision^15–18^. On each trial, participants were presented with 8 circles colored between red and blue, and they had to judge whether the average color was closer to red or to blue. We compared two conditions (Figure 4A). Within a block, one set of trials had circles with high mean evidence (the average color was far away from the midpoint between red and blue) and with high evidence variability, while the other set featured trials with low mean evidence (the average color was close to the midpoint) and low variability (more homogeneous hue across circles). These two conditions were matched in terms of decision accuracy (t(28) = −0.993, *p* = 0.329, 95% CI [-0.080, 0.028], *d* = −0.184, see Figure 4B), but confidence ratings collected after the main experiment confirmed that participants were slightly more confident in the *low* condition (t(28) = −3.151, *p* = 0.004, 95% CI [-5.855, −1.242], *d* = −0.585, see Figure 4C), as found in previous studies with these stimuli^16,17^. We thus expected to find that participants would prioritize the low condition over the high condition. Overall, this effect was not clear at the group level: the proportion of blocks where the low condition set was chosen first was not different from 0.5 (t-test: t(28) = 0.851, *p* = 0.402, 95% CI [-0.026, 0.062], *d* = 0.158, Figure 4D). Crucially, however, when looking at the inter-individual heterogeneity, we found that the expected priority effect towards the low condition was more pronounced for participants expressing greater confidence in the low condition when compared to the high condition (Figure 4E). Given the presence of a clear outlier (who always prioritized the first presented set, irrespectively of the condition), the relation between these two effects was confirmed with a robust correlation analysis (*r* = 0.427, *p* = 0.021). Of note, prioritization of the low condition was not correlated with a performance advantage towards the low condition (*r* = - 0.099, *p* = 0.611). After excluding the outlier, we could also predict the priority effect from the confidence effect in a regression across participants (*β* = 0.015, *t* = 3.399, *p* = 0.002), even after adding the performance difference between the two conditions as a covariate in the regression (*p* = 0.253). In sum, Experiment 5 shows that participants who experienced more confidence in the low condition also tended to prioritize this condition in their task planning, even when task accuracy was equal.

**Figure 4.**
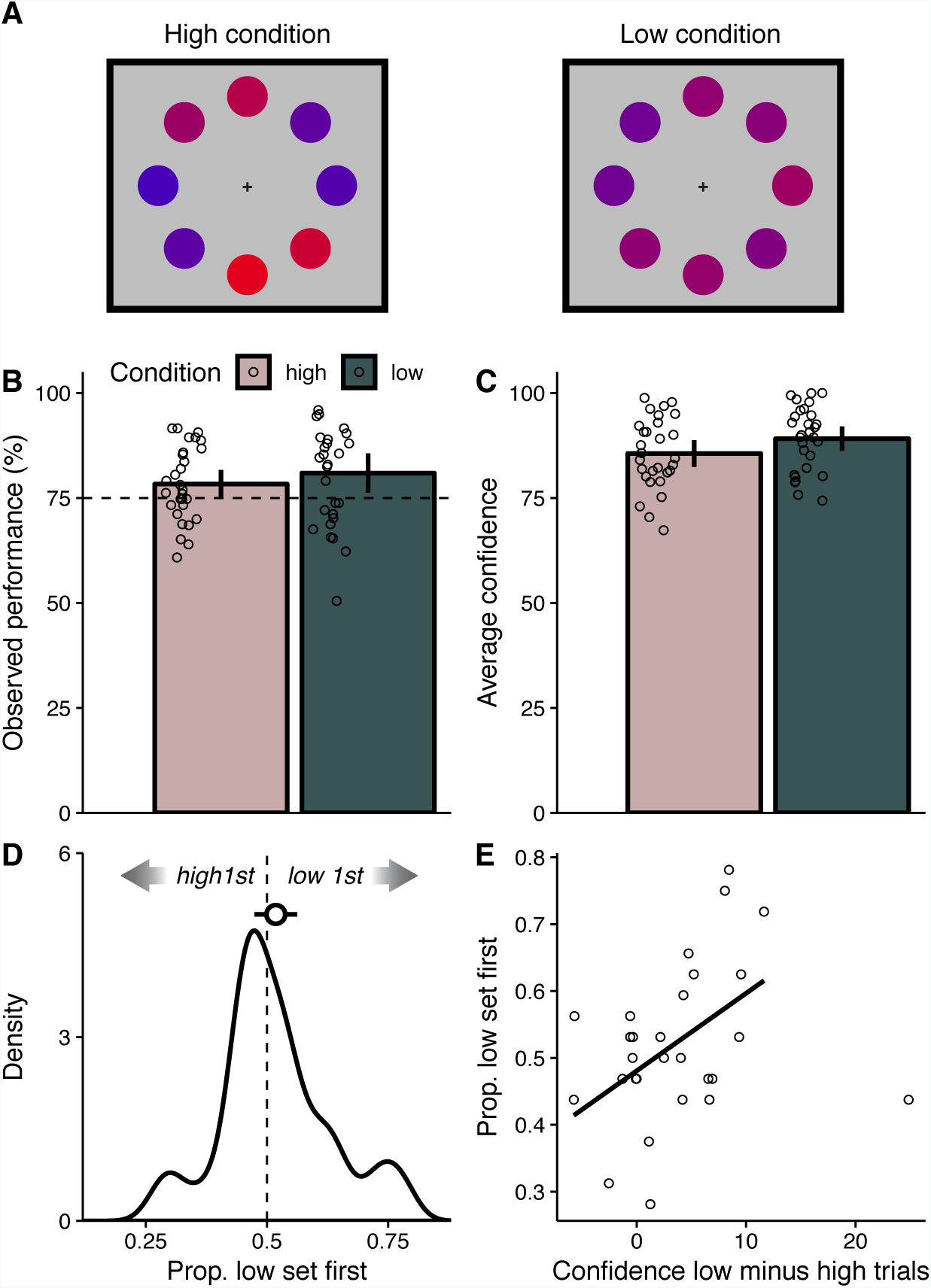
Experiment 5: design and results. **A.** Example stimulus for each of the conditions. The mean evidence generating each sample from the high condition is of high evidence, but across-sample variability is also high. For the low condition, the generative mean is of lower evidence, but variability is also lower. **B.** For all trials of the experimental part, average observed performance as a function of expected performance, both in percentage, split by condition. Bars represent averages across participants, with error bars denoting 95% confidence intervals. Dots represent individual participants. The horizontal dashed line indicates where 75% observed performance would lay. **C.** Average confidence ratings across participants, split by condition. Error bars denote 95% confidence intervals. Dots depict individual participants. **D.** Density of the proportion of blocks where participants chose to complete first the the set whose trials belonged to the low condition. The density is drawn by using each participant’s data. The across-participant average is displayed as a big dot, with error bars showing 95% confidence intervals. The dashed line corresponds to low and high condition sets being chosen first equally often. **E.** Proportion of blocks where participants chose to complete first the the set whose trials belonged to the low condition, as a function of the difference between the average confidence for low condition trials minus the average confidence for high condition trials. Each dot depicts an individual participant. The line depicts a linear fit of the data excluding the outlier (rightmost dot).

Finally, we aimed at evaluating the generalizability of our findings beyond perceptual decisions. For this, we conducted another experiment to test whether priority towards easier problems would also manifest itself when considering simple mental calculation. On each trial, two schematics of mental calculation problems were presented (Figure 5A). One was a priori more difficult than the other, in the sense that it included a multiplication or a division (e.g. “x+y/z”), instead of only additions or subtractions (e.g. “x+y-z”). The problems were otherwise matched in the number of elements and operations they involved. Participants had to click on one of the schematics to reveal the actual problem, which they then had to solve. Immediately after, participants had to solve the non-chosen problem. Although there was no difference in response accuracy between easy and hard problems (*p* = 0.883), the response times for the hard problems were significantly longer (Figure 5B, t(100) = 10.001, *p* < 0.001, 95% CI [3.010, 4.499], *d* = 0.995), proving that they were more demanding. Most participants predominantly chose to solve the easy problem first (Figure 5C, t(100) = 6.146, *p* < 0.001, 95% CI [0.590, 0.676], *d* = 0.612). This experiment thus replicated and extended beyond perceptual tasks our previous findings of an easy-first bias in task prioritization.

**Figure 5.**
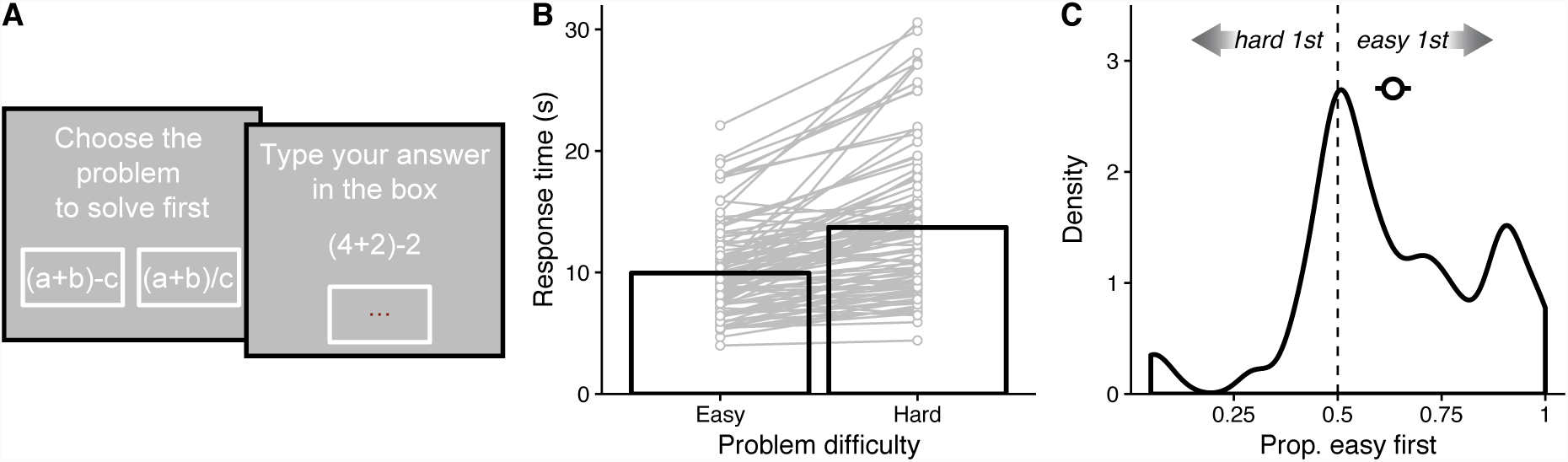
Experiment 6: design and results. **A.** Overview of one trial. Two schematics of simple mental calculation problems were presented, one easier to solve than the other. Participants clicked on the one they wanted to solve first. The actual problem was then revealed, and participants had to type their response. Then they had to do the same with the other problem. **B.** Average response time, in seconds, as a function of the difficulty of the problem. Bars show the average across participants, while dots connected by lines represent each individual participant. **C.** Density of the proportion of blocks where participants chose to respond first to the easy problem. The density is drawn by using the proportion for each participant. The across-participant average is displayed as a big dot with error bars showing 95% confidence intervals. The dashed line corresponds to easy and hard problems being chosen equally often.

To summarize, we found that, when having to prioritize responses or tasks, participants exhibited a systematic bias towards engaging first in the response or the task in which they had greater confidence. We replicated this finding in 6 experiments. Converging evidence came from different measures, using task difficulty and observed accuracy as proxies for confidence, but also using actual confidence ratings, and even exploring situations where performance and confidence were dissociated. In terms of generalizability, we demonstrated that this effect was present in both situations involving retrospective confidence (experiments 1-3) and prospective confidence (experiments 4-6), and both situations involving perceptual and non-perceptual tasks, such as mental calculation.

Situations in which we face multiple tasks present a huge diversity, and task prioritization is likely determined by a varied set of factors, most of which were not manipulated in our design. For instance, prioritization can be driven by personal interest and the familiarity with the task^19^. We did also not manipulate the duration of each task, although there is evidence from real-life situations where people tended to approach first the shortest tasks^20^. Although in our experiments the two tasks were presented simultaneously, previous research has shown that task prioritization also depends on the time at which task-relevant information is presented^21^. We therefore only claim that confidence is one among many forces driving task prioritization. In our data, we find a large inter-individual variability in the general tendency to prioritize the color or the letter task (Figure 2A, see also supplementary information 6.1.), and we found that the locations of the different choice options on the screen could also bias task priority (see supplementary information 6.2.). Whether this variability is due to strategies or heuristics explicitly chosen by participants, or to differences in their perceptual or decision processes, remains to be investigated. Another factor that deserves more scrutiny is the reward associated with each task, which should affect task priority. In our experiments, we purposefully avoided any difference in reward between both tasks within a trial. We also did not instruct or incentivize participants to prioritize the more confident or the more accurate of the two decisions. In sum, nothing in the instructions or the reward structure pushed participants to prioritize one task over the other. Some of the aforementioned features could be manipulated in future studies, and whether humans change their priorities under such manipulations is an open empirical question.

Another important issue for future research is whether this strategy of approaching the easy task first is adaptive. A priori, reducing the mental workload before tackling more difficult tasks could be beneficial. In our data, however, we found no clear evidence that addressing the easy task first led to better performance. Overall, participants were neither better nor worse when responding first to the easy task than when responding first to the hard task (see supplementary information 7.1.). To further investigate this issue, we conducted an additional experiment comparing a free response order condition (which replicated Experiment 2) to an imposed order condition in which the choice for the hard task was required either first or second (see supplementary information 7.2.). We found that participants’ performances were similar between these two conditions, suggesting that, in our paradigm, the confidence-driven prioritization does not lead to an advantage in task performance. In other words, while we cannot claim that there is no advantage behind confidence-driven prioritization, our manipulation could not find it.

Our study highlights a new way in which confidence actively shapes our behavior. It therefore contributes to a growing literature showing how confidence is not a mere commentary on performance but actually affects individual or collective decisions. Previous research showed how, when agents interact, the confidence with which an advice is expressed affects the use of this information by others^22–24^ and how well the group will perform^25^. At the individual level, confidence may serve as a teaching signal when feedback is unavailable^26–29^. It also influences the amount of resources we engage in a task^17,30^. Here we show that confidence does not only determine how we do a task, but also when we plan to do it.

Understanding how people decide to perform one task before another may have practical consequences for the management of individuals and organizations. This motivated some studies that focused on different applied scenarios. Some of this previous work already found that individuals address easier tasks first when they can decide their task schedule. For instance, students appear to prioritize easier course assignments over difficult ones^31^, and the expected easiness of finding the relevant information affects their prioritization of a web-search task over another^19^. Physicians at an emergency department do prioritize easy patient cases, especially under high workload contexts^7^. The present study makes several unique contributions to this research topic: we extend this result to situations involving immediate decisions, we link the easy-first bias to confidence, and we offer a strict control over performance in the task via our psychophysical procedures. Finally, although we did not find a clear consequence from the easy-first bias on performance, previous research has shown that in another context this strategy was associated with both short-term benefits and long-term costs. Indeed, prioritizing easy cases allowed physicians to treat more patients per unit of time, but across individuals it was associated with a lower productivity and less revenue for the hospital^7^. Understanding whether and how similar short-term benefits and long-term costs may arise in other contexts, and in particular in situations where individuals have to perform multiple perceptuo-motor tasks at the same time, constitutes an exciting topic for future research.

## Methods

### Participants

Participants (Exp1=28, Exp2=20; Exp3=30; Exp4=31; Exp5=29; Exp6=101) were healthy adults recruited from the Laboratoire d’Économie Experimentale de Paris. They reported normal or corrected-to-normal vision, and provided informed consent. Participation was compensated based on performance and duration, at an average rate of 12€ per hour. The Paris School of Economics ethics committee approved the experimental procedures after determining that they complied with all the ethical regulations. Participants were tested in sessions of up to 20 individuals. The number of participants in each experiment depended on the number of people showing up for that particular experiment. We aimed at a minimum of 20 participants in each experiment.

### Apparatus

Experiments were run using MATLAB Psychtoolbox^32^ on a computer screen (17’, 1024×768 pixel resolution, 60 Hz refresh rate) viewed from approximately 60 cm. Stimuli appeared on a grey background.

### Perceptual task (experiments 1-4)

For experiments 1-4, the task consisted in reporting the dominant letter and the dominant color of an array of 80 letters (Xs or Os), colored blue or orange (see Figure 1A). Each trial started with a 500 ms fixation cross, followed by a 300 ms blank period, and then by the stimulus, which was presented for 1 s. Each letter, in Arial font, occupied approximately 0.5º (degrees of visual angle). Elements were presented within a 10º wide imaginary square. These stimuli were based on the code available at https://github.com/DobyRahnev/Confidence-leak and used in a previous publication^33^.

For Experiments 1-3, after the stimuli disappeared, the response screen was presented, which consisted in 4 square boxes that contained a blue square, an orange square, a white O and a white X (see Figure 1A). The 4 boxes were randomly arranged in a 2×2 grid on each trial. Participants gave their response by clicking on a color box and a letter box, in their preferred order. A first response on one dimension (e.g. clicking on the X box) made the corresponding boxes disappear, leaving only the boxes for the other dimension (e.g. the orange box and the blue box), which required a second response. Participants completed 4 parts of 4 blocks each, with 25 trials per block. Between block and block, participants could rest for 15 seconds, and at the end of each part they could take self-timed breaks.

### Manipulation of task difficulty (experiments 1-4)

For each perceptual task, difficulty was manipulated by changing the proportion of the dominant over the non-dominant feature. After a short training, we used psychophysical staircases to estimate, separately for each task, the proportion of items that should be used for responses to be 90% correct (*easy* condition) or 60% correct (*hard* condition). Details of the training and staircase procedure are presented in supplementary information 1.1. We then used these parameters in the main experimental part to manipulate difficulty on the letter (easy vs. hard) and color (easy vs. hard) tasks, orthogonally within each participant, in a 2×2 factorial design. For experiments 1-3, we collected 100 trials for each of the 4 conditions, with trials of all conditions intermixed within blocks.

### Experiment 1: confidence ratings

In Experiment 1, after participants made their two choices, they had to rate their confidence on each of them. Two confidence scales were simultaneously presented, one horizontal for the color task and one vertical for the letter task (see Figure 1A). Each scale featured the two response categories at its ends (which corresponded to 100% confidence) and an empty region at its center (near which confidence was at 50%). The two choices participants had made on that trial were surrounded by a yellow circle. Participants gave their confidence levels on both tasks by clicking on the corresponding scales, and then validated the ratings by clicking at the center of the screen. Participants were told that maximum uncertainty (no preference between their response and the other option within the same task) corresponded to a 50% confidence, and absolute certainty on the choice to a 100% confidence.

### Experiment 2: choice-only

Trials in Experiment 2 closely resembled those in Experiment 1, except for the fact that no confidence had to be rated. Thus, the trial finished after participants made a choice for each task on the response screen.

### Experiment 3: varying delays

In Experiment 3, we manipulated the time between stimulus offset and the presentation of the response screen, which was set to 0, 2 or 4 seconds (counterbalanced across trials).Considering the response times for our previous experiments (see supplementary information 3.4.), 4s ensured that participants had reached both decisions by the time the response screen was presented.

### Experiment 4: anticipated difficulty in prospective confidence

In Experiment 4, participants only responded to one task on each trial. Therefore, the response screen uniquely contained the two relevant boxes. The main part was organized in 32 blocks. All blocks included a familiarization and a test phase (Figure 3A). During the familiarization phase, participants completed two sets of 6 trials. Each set was associated to a single task (color or letter), a difficulty level (easy or hard), and an animal name (randomly chosen without replacement from a predefined list of common names). Based on staircases ran before the main phase, the proportion for the task-relevant dimension was experimentally controlled, such that for each block, one set contained difficult trials and the other easy trials (for the non-relevant dimension, the proportion was always set at 50%). This manipulation was not revealed to participants but they could learn the difficulty of each set by performing the task during the familiarization phase. After the familiarization, the test phase started. Participants saw a screen with two horizontally aligned boxes presenting (in a random position) the animal names of the two sets just completed. 4 trials of each set had to be completed, with reward in that block depending on performance in that part. Participants decided the order in which to complete the test trials of each set by clicking on the name of the set they desired to complete first. We used animal names to avoid any obvious order bias (set A vs B, or 1 vs 2) for this decision (for the complete list of set names, see supplementary information 9.1.). Across blocks, we manipulated factorially the task (color/letter) used for the first set in the block, the task (color/letter) used for the second set, and the order of the difficulty levels used for the two sets (easy first vs. hard first). This design resulted in 8 conditions, presented in a randomized order as a sequence 4 times per participant.

### Experiment 5: dissociation between performance and confidence

For Experiment 5, participants had to categorize the average color of a set of 8 circles as more red or blue. Each trial started with a black fixation cross presented for 400 ms, followed by 8 circles presented simultaneously for 200 ms. Each circle was uniformly colored in RGB space as [255*t, 0, 255*(1-t)] with a color parameter t taking values between 0 (full blue) and 1 (full red). The circles, each with an area of 2500 pixels, were regularly spaced on an imaginary circle 120 pixels from fixation. After the stimulus disappeared, the fixation cross turned white and participants had to make a response. For all parts of the experiment, within each block of trials, red was the dominant component (and thus the correct response) for 50% of the trials, and blue for the other 50%. Participants had to use the E and R keys of the keyboard to indicate whether the average color was closer to red or to blue. The mapping between each of the two keys and each of the two color responses was randomized across participants.

On each trial, the color parameter t for each circle was sampled from a standard normal distribution, then multiplied with the standard deviation of the corresponding condition (high: SD= 0.20, low: SD=0.05) and then added to the mean of the corresponding condition, which was determined with a staircase procedure at the beginning of the experiment (see Supplementary Material 1 for details).

The main part of this experiment greatly resembled that of Experiment 4. On each of its 32 blocks, participants were introduced to two sets of trials, each with an animal name: for one set, the generative mean was high, but so was the SD (high condition), whereas for the other both mean and SD were low (low condition). After a familiarization phase where 6 trials of each set were completed, participants had to complete 4 more trials of each set, with these test trials determining payoff at the end of the experiment. However, they could choose the order in which to complete them by clicking on the name of the set they wanted to start with. Apart from the nature of the stimulus, the main difference with respect to Experiment 4 was that the expected difficulty of both sets was always 75%, with the difference between sets laying on the mean and variability of the circles that were generated on each trial. For each participant, we achieved the values leading to a 75% expected performance through an initial staircase procedure for each condition. For more details, see supplementary information 8.

After the main part of this experiment, participants completed a part where 8 blocks of 24 trials were presented. Each trial could have the mean and SD values previously used for the high condition, or those used for the low condition. These two types of trials were interleaved within each block, with each type appearing the same number of times. The crucial feature of this final part was that, after their response, confidence had to be reported. Confidence was given by clicking on a vertical scale that, as in Experiment 1, ranged from 50 to 100. Confidence was only required in this part, and not before, to avoid biasing the participants’ priority choices in the previous part.

### Experiment 6: anticipated difficulty in a mental calculation task

The stimuli used in Experiment 6 were different from those used in the previous experiments. On each trial, two horizontally aligned boxes appeared (see Figure 5A), each containing a simple formula (e.g. “a+2b”). Our formulas involved letters connected by basic operators (+, -, *, /) and were sometimes nested by brackets. Both problems were similar except for the operators connecting their respective letters. One of the problems (easy) contained additions and subtractions, while the other (hard) contained multiplications and divisions. The easy and hard box were randomly positioned on the left or the right of the screen on every trial. Participants had to solve both problems in their prefered order. By clicking on one box, the letters of the formula in that box were replaced by one-digit integers. Participants then typed their response, which appeared inside a box below the problem. They could change it as they wanted before pressing the ENTER key to validate it. Then, the non-clicked on problem appeared for participants to give their response through the same method. Each participant completed the same 20 pairs of questions (listed in supplementary information 9.2.), but in a random order. The initial presentations of formulae ensured participants could not try to solve the problems before making a choice.

### Payoff

Performance on the main part of the experiments determined the final amount of money participants won. In experiments 2 and 3, for each of the 16 blocks, a random trial was chosen. Within that trial, the choice for either color or letter was also randomly picked. If that choice had been correct, the participant received 1 €, and 0 € otherwise. This payoff method ensured participants kept a stable performance and did not relax their attention or automatized their responses during some trials.

In Experiment 1, the payoff system was slightly different. Once a trial and a choice had been selected within a block, a number was selected from a uniform distribution between 0 and 100. If the number was lower than the confidence given for that choice, the payoff depended on performance. If the number was higher, another random number was sampled from a uniform distribution between 0 and 100. If this number was smaller than the random number previously drawn, 1€ was given, and 0€ otherwise. This system has previously^34^ been adopted to ensure that participants try to accurately rate their confidence, since their payoff /depends on it. On top of this, participants were awarded a fixed 3€ just for the added length reporting confidence gave to the experiment.

In Experiment 4, payoff was similar to Experiment 2, but given the more numerous and shorter blocks, each of them could be rewarded with only 0.5€. The rest between blocks was also shorter, with a duration of 10 s.

Experiment 5 had a payoff like Experiment 4, but we also added an extra compensation for the final trials where confidence had to be reported. For each of the 8 blocks in which trials were divided, participants could receive 0.5€ in a procedure like that used in Experiment 1, where reward could depend on performance or on confidence.

Experiment 6 had a payoff system which mimicked that of the previous experiments. At the end of the experiment, two trials were chosen randomly. For each of these selected trials, the answer to one of the problems was picked. If the answer was correct, participants received 1€, and nothing otherwise.

Before starting each experiment, participants were carefully informed about the procedure. The payoff structure and its objective were made clear, and the fact that reward did not depend on their response times was also pointed out.

### Statistics

All reported t-tests and binomial exact tests are two-tailed. All ANOVA are repeated measures. For t-tests, effect size is reported by using Cohen’s *d*_*z*_, written here as *d*, which was calculated using Rosenthal’s formula^35^: the *t* statistic was simply divided by the square root of the sample size. For ANOVA, effect size is reported by using partial eta squared 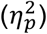. For each factor, this was calculated by taking the sums of squares of the condition, and dividing it by the sums of squares of the condition plus the sums of squares of the error^36^. Logistic regressions were performed with R’s function glmer, by using maximum likelihood and a logit link function.

### Code availability

The code used to produce the analyses and the figures that appear on this study are available from the corresponding author upon reasonable request.

## Acknowledgements

This work was supported by the Agence Nationale de la Recherche [grant ANR-16-CE28-0002 to VdG]. We thank Maxim Frolov, Audrey Azerot and Quentin Cavalan for helping in data collection. We thank Mahiko Konishi, Jean-Christophe Vergnaud, Jérôme Sackur, Bruno Berberian and Pascal Mamassian for discussions. We thank Dobromir Rahnev for making available his experiments’ code.

## Supplementary information

### 1. Training and calibration for experiments 1 to 4

#### 1.1. Procedures

Participants started the experiment with 10 training trials for each task in isolation. During the color training, all the 80 letter elements were either O or X. The color task also started at its easiest, with all the 80 elements being of one color, but it became more difficult at the rate of one element per trial. The predominant color and letter were determined randomly on every trial. After the stimuli, only the two boxes for the color task appeared, and the participant gave a response only for that task. A similar procedure was followed for the letter training block, but with all the elements being one color, and the letter task getting more difficult. The order in which these two training blocks were presented was counterbalanced across participants.

After the training, participants were told that the real experiment was about to start, composed by 5 parts of 4 blocks each. In the first part, the proportion of elements was manipulated with a staircase procedure that would allow the presentation of performance-adjusted easy and hard trials in the subsequent parts. This procedure was not disclosed to participants, to avoid any temptation to manipulate it. The staircase followed a one-up/one-down procedure with unequal up and down steps (see e.g. García-Pérez, 1998). Specifically, the initial number of elements in the dominant category was set to 64 out of 80 elements (80%-20%) for both tasks, and, it was decreased by one element after a correct response, and increased by four elements after an error. We then used the 100 trials of that first part to estimate for each task a psychometric function, so as to generate parameters for the following parts. For the color task, this function represented the probability of choosing blue as a function of the number of blue elements, and was fit with a cumulative Gaussian; the same for the probability of choosing O for the letter task. To calculate the number of elements corresponding to a 90% performance (our easy condition) in the color task, we used 40 plus or minus the semi-difference between the number of blue elements for which the psychometric curve predicted a 90% and 10% of blue choices, rounded to its nearest integer. Note that by doing so, we assumed that in the main part participants would not be biased towards one response category. Similar procedures were carried out to obtain parameters for the 60% performance (hard condition), and for both difficulties of the letter task. As can be seen in the results section of the main text and in Figure S1, observed performance in the main experiment closely matched the targeted performance.

In Experiment 4, since each trial only required responding to one task, previous to the main part two staircases were run sequentially, one for the color task and one for the letter task, with a randomized order across participants. Each of them had 100 trials, and the elements of the non-relevant task were kept stable at a proportion of 40/40. These trials were presented to participants as a training part, before the main experiment.

#### 1.2. Mean and SD of stimulus elements

The following table indicates the number of elements in the dominant category, in the easy condition and in the hard condition, for the color task and the letter task. Means and standard deviations across participants are reported. These parameters were estimated using a staircase procedure for each task (see previous section) and used in the main part of experiments 1 to 4. However, the color and letter staircase procedures were run in parallel for experiments 1 to 3, and one after the other for Experiment 4. The total number of elements in the display was always 80, and the number of elements in the non-dominant category was adjusted accordingly.

|  | Color task |  |  |  | Letter task |  |  |  |
| --- | --- | --- | --- | --- | --- | --- | --- | --- |
|  | easy |  | hard |  | easy |  | hard |  |
| Exp # | M | SD | M | SD | M | SD | M | SD |
| 1 | 48.3 | 2.62 | 41.8 | 0.585 | 62.1 | 7.58 | 44.3 | 1.47 |
| 2 | 47.7 | 3.06 | 41.5 | 0.761 | 56.7 | 6.21 | 43.3 | 1.26 |
| 3 | 47.9 | 2.37 | 41.5 | 0.509 | 56.4 | 4.84 | 43.2 | 0.971 |
| 4 | 45.8 | 1.54 | 41.1 | 0.301 | 55.4 | 3.91 | 43.2 | 0.860 |

#### 1.3. Expected and observed performance

**Figure S1.**
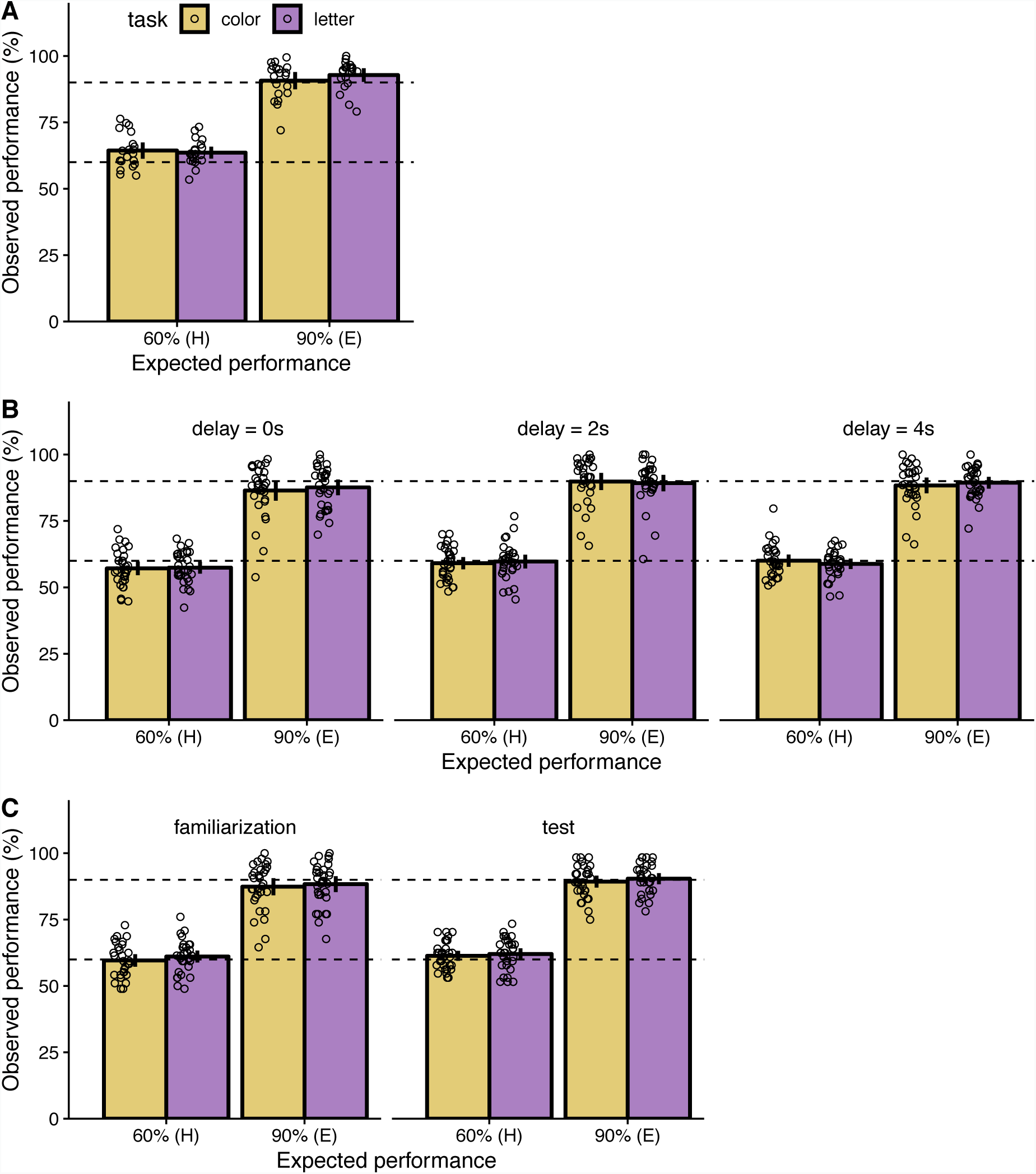
Expected and observed performance for experiments 2 to 4. **A.** For Experiment 2, observed performance as a function of expected performance, both in percentage, split by task type. Bars represent averages across participants, with error bars denoting 95% confidence intervals. Dots represent individual participants. Top and bottom dashed lines help indicate where 90% and 60% observed performance would lay, respectively. **B.** Same as A, but separately for each time delay of Experiment 3. **C.** Same as A, but separately for the familiarization and test phase of Experiment 4.

### 2. Training and calibration for Experiment 5

Before the main part and the confidence trials, two other parts were run.

The experiment started with 20 easy training trials that familiarized participants with the structure of each trial. After each response, feedback on their performance was given by displaying the message ‘CORRECT’ or ‘ERROR’.

After the training trials, participants were told that they were going to do a second, longer training session. In fact, in this session the generative mean of the stimulus parameter t was updated according to a staircase procedure, to obtain 75% accuracy in both the high condition and the low condition. We used a one-up/one-down procedure with unequal up and down steps. The starting value for the mean was 0.75, and it was reduced by 0.0078 after a correct response and increased by 0.032 after an error. This mean parameter determined the signal strength towards red, and on a random half of the trials we used 1-t to obtain blue trials. This staircase procedure was conducted separately for the high condition (SD=0.2) and low condition (SD=0.05) in two interleaved staircases of 100 trials each. When the staircase phase was finished, we estimated for each condition the value leading to an expected performance of 75%. We did so as in previous experiments. Importantly, during this part feedback on performance was still given after each trial in order for participants to learn where the midpoint between red and blue laid. Feedback was removed for the main part and for the final trials where confidence reports were required.

### 3. Supplementary analyses for Experiment 1

#### 3.1. Confidence in the first choice vs. in the second choice

In the main text, Figure 1D shows the distribution of confidence pooled across all trial types, but also pooled across same-difficulty trial types (EE and HH combined). In order to provide more detailed information, Figure S2 shows the information for each trial type, with Figure S2A showing the data for the whole dataset, Figure S2B showing the data only those trials where both choices were correct, and Figure S2C showing the data only for those trials where both choices were errors.

**Figure S2.**
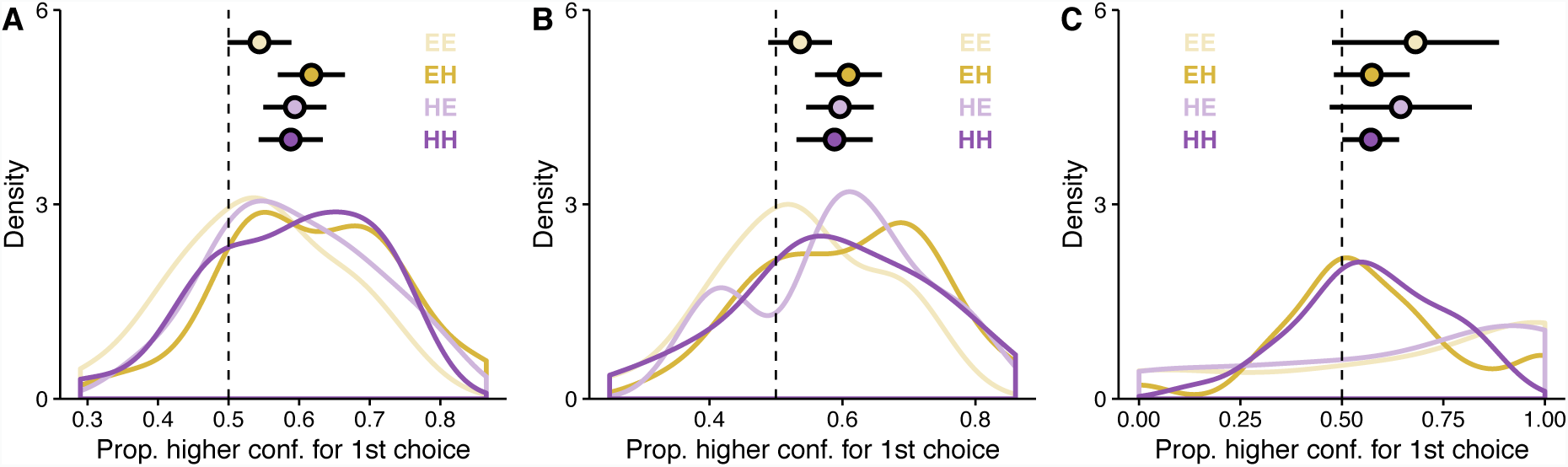
Confidence in the first vs. in the second choice. **A.** For Experiment 1, density of the proportion of trials where the first choice had a higher confidence rating than the second choice, split by trial type. Densities are drawn by using each participant’s data. Across-participant averages are displayed as big dots with error bars showing 95% confidence intervals. The dashed line corresponds to equal confidence ratings for first and second choices. **B.** Same as A, but taking only trials in which both choices were correct. **C.** Same as A, but taking only trials in which both choices were errors.

#### 3.2. Confidence as a function of difficulty and accuracy

Except for Experiment 1, we did not ask participants for confidence ratings. Thus, we had to rely instead on different proxies for confidence to find support for our priority effect. We used expected difficulty and response accuracy as proxies for confidence. These measures have repeatedly been found to correlate with confidence (Kepecs, Uchida, Zariwala, & Mainen, 2008; Peirce & Jastrow, 1885; Sanders, Hangya, & Kepecs, 2016). We first checked that such links were also present in Experiment 1. To do so, we fitted a linear mixed model (LMM) with average confidence as dependent variable, and task type (color vs. letter), accuracy (correct vs. error) and expected difficulty (easy vs. hard) as fixed effects. Participants were treated as random effects. We chose an LMM over an ANOVA due to the big difference between the number of correct and error trials. Two participants were excluded due to the absence of any error trial when the tasks were easy. The fixed effects of expected difficulty (*t*(20767.359) = 5.697, *p* < 0.001) and accuracy (*t*(20767.635) = 20.401, *p* < 0.001) were significant, but also their interaction (*t*(20767.331) = −13.193, *p* < 0.001). As Figure S3A illustrates, the average confidence ratings were higher for easy than for hard stimuli when the choice made by participants had been correct, and lower for easy than for hard stimuli when errors are considered. The fixed effect of task type was not significant (*p* = 0.121), and it produced no significant interaction with expected difficulty (*p* = 0.379), but it did with accuracy (*t*(20767.444) = 3.427, *p* < 0.001), and there was a triple interaction among all the fixed effects (*t*(20767.305) = −2.352, *p* = 0.019). This pattern of results replicates classic findings and reaffirms the validity of the measures we used for the analyses in Experiment 2.

#### 3.3. Priority effects from task difficulty in Experiment 1

After establishing that task difficulty was associated with confidence in our setup, we could evaluate their relationship to priority, as we did in our analyses of experiments 2 and 3 (Figure 2). We report here the same analyses applied to Experiment 1. We can mimic Figure 2A and plot the proportion of choosing color first across the different trial types (Figure S3B). Individual task biases toward the color or the letter task can be seen, as well as the difference between EH and HE trials. This difference can be better appreciated in Figure S3C (that resembles Figure 2B). The color task was first responded to on 66% of EH trials and on 50% of HE trials. The difference was significant across participants (t(27) = 6.231, *p* < 0.001, 95% CI [0.107, 0.212], *d* = 1.178).

**Figure S3.**
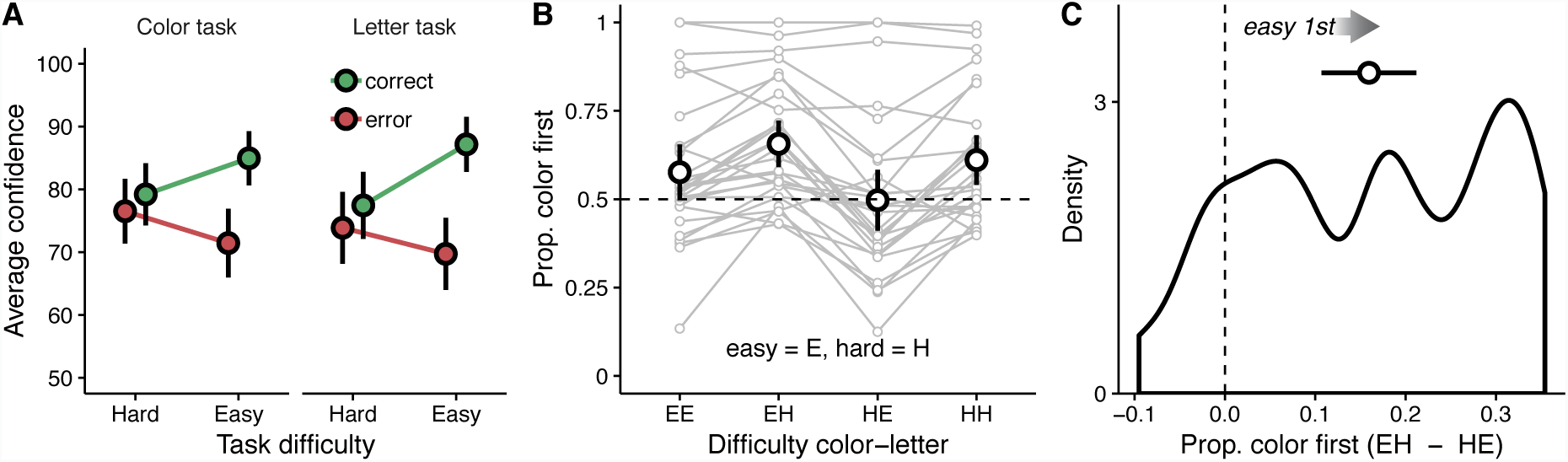
Experiment 1: supplementary data. **A.** Average confidence ratings across participants, as a function of task difficulty, and split between correct and error trials (color-coded) and by task type (different panels). Error bars denote 95% confidence intervals. **B.** Proportion of trials where the color task was first responded to, as a function of the type of trial (EE = easy color, easy letter; EH = easy color, hard letter; HE = hard color, easy letter; HH = hard color, hard letter). Each set of four dots connected by lines represents one participant. Bigger dots represent average proportions across participants, and error bars denote 95% confidence intervals. The dashed line corresponds to no preference towards responding first to the color or the letter task. **C.** Density of the difference between proportion color task chosen first for EH trials and proportion color task chosen first for HE trials. The density is drawn by using the difference of proportions for each participant. The across-participant average is displayed as a big dot with error bars showing 95% confidence intervals. The dashed line corresponds to no difference between EH and HE trials.

#### 3.4. Response times

Experiment 3 implemented a variable delay between the removal of the perceptual stimuli and the presentation of the response screen. The values the delay could acquire were decided based on the typical response times in our previous experiments. Figure S4 shows the distribution of response times for the first and second choices in Experiment 1. Response times correspond to the time between the presentation of the response screen until a click was made on a response square (first choice), or to the time between this first click and a second click corresponding to the other task (second choice). Longer response times for the first choice are most likely due to the initial decision of which task to respond to first, plus the initial identification of how each element was mapped on the screen. For the two non-zero delay values in Experiment 3, we selected 2s, and 4s. The first value was longer than the response time for most choices, and the second was big enough to assume that the decision for the hard task should already have been reached.

**Figure S4.**
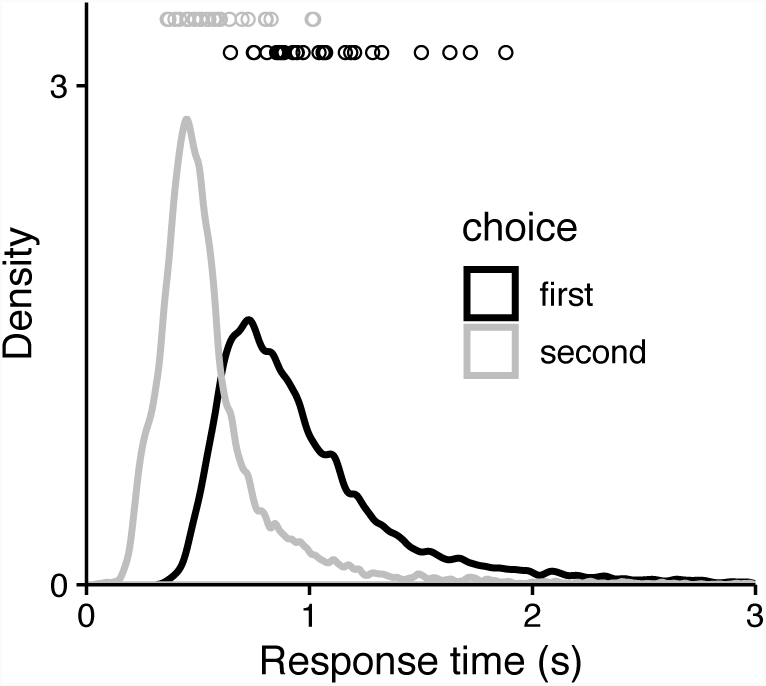
Response times density of the first and second choice in Experiment 1. Densities are drawn by using the whole distribution of trials for all participants. Participant averages are depicted as single dots.

### 4. Logistic regression tables for Experiment 1

The next tables contain the fixed effects of the logistic regressions that were ran for Experiment 1, as well as the comparison between models.

In all models, the dependent variable was always the probability of responding to the color task first, and participant id was included as a random effect.

In what follows, we use the following abbreviations for the predictors:

- DiffExpPerf: difference in expected performance (color minus letter).
- DiffActAcc: difference in accuracy (color minus letter).
- DiffConf: difference in reported confidence (color minus letter).

Analysis 1: Logistic regression using the full model

All data, full model, logistic regression

| | $\beta$ estimate | Standard error | z value | $p$ |
| --- | --- | --- | --- | --- |
| Intercept | 0.616 | 0.241 | 2.554 | 0.011 |
| DiffExpPerf | 0.003 | 0.001 | 2.910 | 0.004 |
| DiffActAcc | 0.002 | 0.000 | 4.936 | <0.001 |
| DiffConf | 0.045 | 0.002 | 29.474 | < 0.001 |

Analysis 2: Comparison with a ‘partial model’ without the confidence predictor

All data, partial model, logistic regression

Analysis 2: Comparison with a 'partial model' without the confidence predictor All data, partial model, logistic regression
| | $\beta$ estimate | Standard error | z value | $p$ |
| --- | --- | --- | --- | --- |
| Intercept | 0.574 | 0.244 | 2.353 | 0.019 |
| DiffExpPerf | 0.009 | 0.001 | 8.886 | < 0.001 |
| DiffActAcc | 0.004 | 0.000 | 9.742 | < 0.001 |

All data, model comparison between partial and full model

| Model | Deg.freedom | AIC | BIC | logLik | deviance |
| --- | --- | --- | --- | --- | --- |
| partial | 4 | 13296.734 | 13326.029 | -6644.367 | 13288.734 |
| full | 5 | 12177.248 | 12213.866 | -6083.624 | 12167.248 |

$\chi^2(1) = 1121.4864, p < 0.001$

Analysis 3: Two-step regression approach

Here, we used the predicted values of the ‘partial model’ as an offset in another regression, in which priority was predicted by the ‘residual confidence’ (ResidConf), that is the confidence (DiffConf) left unexplained by accuracy (DiffActAcc) and expected performance (DiffExpPerf).

| | $\beta$ estimate | Standard error | z value | $p$ |
| --- | --- | --- | --- | --- |
| Intercept | 0.026 | 0.022 | 1.191 | 0.234 |
| ResidConf | 0.043 | 0.001 | 26.700 | < 0.001 |

We replicated these analyses and obtained similar results if we only included trials with different difficulty levels for the two tasks (EH and HE trials), to ensure that our results hold when difficulty varies across the two tasks.

Analysis 1bis: Logistic regression using the full model

Different-difficulty data, full model, logistic regression

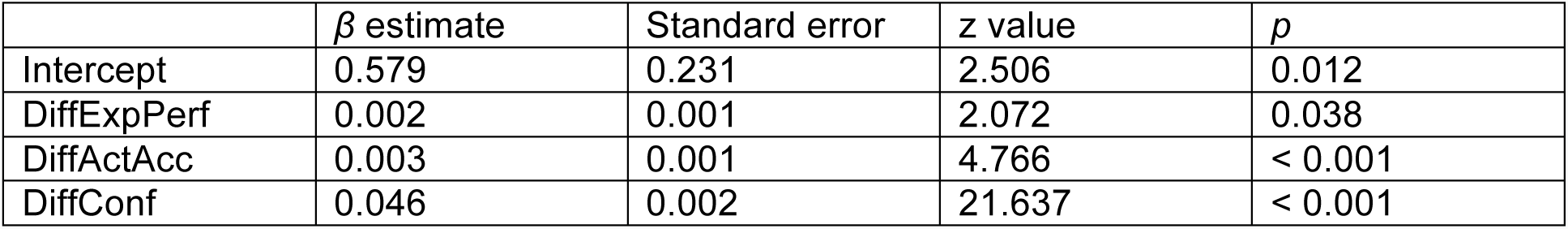

| | $\beta$ estimate | Standard error | z value | $p$ |
| --- | --- | --- | --- | --- |
| Intercept | 0.579 | 0.231 | 2.506 | 0.012 |
| DiffExpPerf | 0.002 | 0.001 | 2.072 | 0.038 |
| DiffActAcc | 0.003 | 0.001 | 4.766 | < 0.001 |
| DiffConf | 0.046 | 0.002 | 21.637 | < 0.001 |

Analysis 2bis: Comparison with a “partial model” without the confidence predictor

Different-difficulty data, partial model, logistic regression

4 Analysis 2bis: Comparison with a “partial model” without the confidence predictor 5 Different-difficulty data, partial model, logistic regression
| | $\beta$ estimate | Standard error | z value | $p$ |
| --- | --- | --- | --- | --- |
| Intercept | 0.520 | 0.234 | 2.222 | 0.026 |
| DiffExpPerf | 0.009 | 0.001 | 7.940 | < 0.001 |
| DiffActAcc | 0.004 | 0.000 | 8.248 | < 0.001 |

Different-difficulty data, model comparison between partial and full model

7 Different-difficulty data, model comparison between partial and full model
| Model | Deg. freedom | AIC | BIC | logLik | deviance |
| --- | --- | --- | --- | --- | --- |
| partial | 4 | 6703.503 | 6703.056 | -3347.751 | 6695.503 |
| full | 5 | 6097.282 | 6130.473 | -3043.641 | 6087.282 |

$\chi^2(1) = 608.221, p < 0.001$

Analysis 3bis: Two-step regression approach

This analysis mimics, with the different-difficulty data, what is performed in analysis 3.

| | $\beta$ estimate | Standard error | z value | $p$ |
| --- | --- | --- | --- | --- |
| Intercept | 0.023 | 0.031 | 0.750 | 0.453 |
| ResidConf | 0.045 | 0.002 | 21.836 | < 0.001 |

### 5. Difference in observed accuracy between first and second choice, for experiments 1 to 3

Apart from confidence ratings and expected accuracy, another validation of the link between priority and confidence could come from data on observed response accuracy. Indeed, since accuracy correlates with confidence in our experimental conditions (as found in our first study), the confidence-priority link could translate into an accuracy-priority link.

For our first three experiments, we looked at whether, within a trial, first choices were on average more accurate than second choices. We found evidence for that in all three experiments (Experiment 1: t(27) = 5.319, *p* < 0.001, 95% CI [0.041, 0.092], *d* = 1.005; Experiment 2: t(19) = 2.969, *p* = 0.008, 95% CI [0.011, 0.066], *d* = 0.664; Experiment 3: t(29) = 2.880, *p* = 0.007, 95% CI [0.007, 0.041], *d* = 0.526).

We also checked if these results held when only including trials with equal difficulty levels (EE and HH) in the analysis. We used this subset of our data to guarantee that this measure was independent from our previous analysis that linked priority to task difficulty by comparing EH and HE trial types. In the same-difficulty case, the comparison of accuracy between the first and second choice was significant for Experiment 1 (t(27) = 2.698, *p* = 0.012, 95% CI [0.007, 0.054], *d* = 0.510) and Experiment 2 (t(19) = 2.097, *p* = 0.049, 95% CI [0.000, 0.060], *d* = 0.469), while it was only a non-significant trend for Experiment 3 (t(29) = 1.856, *p* = 0.074, 95% CI [-0.002, 0.035], *d* = 0.339). The densities of the accuracy differences can be seen in Figure S5.

**Figure S5.**
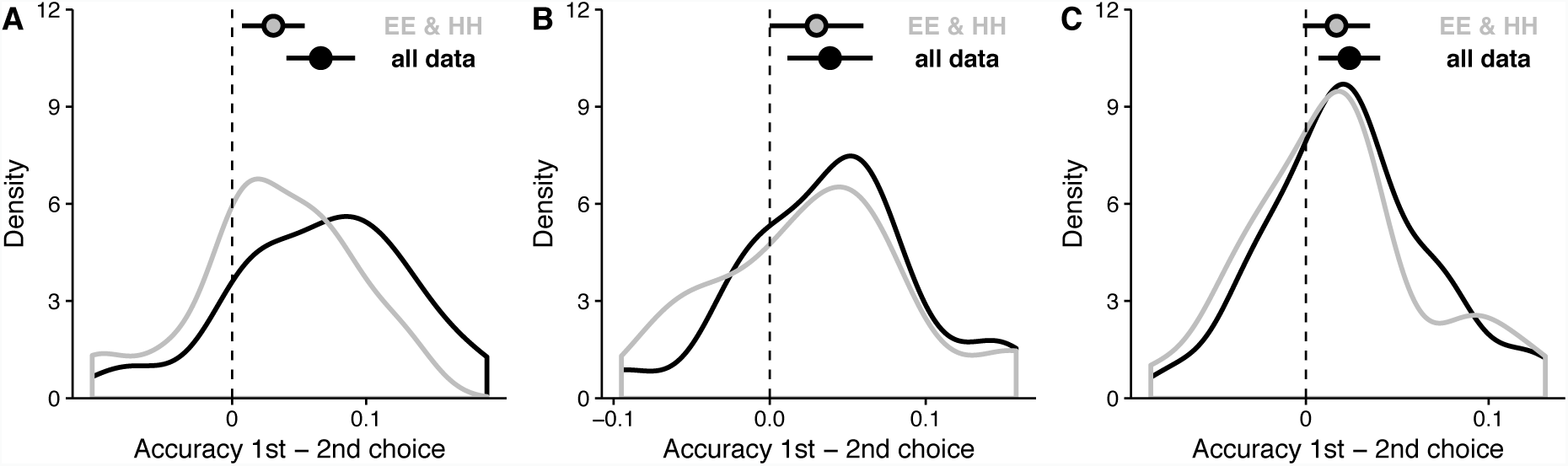
**A.** For Experiment 1, proportion of accuracy on the first choice minus proportion of accuracy on the second choice, displayed as a density across participants. The black density corresponds to the whole dataset, the grey one only to EE and HH trial types. Across-participant averages are displayed as big dots with error bars showing 95% confidence intervals. The dashed line corresponds to no difference in accuracy between first and second choices. **B.** Same as A, but for Experiment 2. **C.** Same as A, but for Experiment 3.

Although we would like to link these results to our analysis relating confidence to priority, there could be however other possible explanations for the fact that accuracy in the first choice is higher than accuracy in the second choice. For instance, memory limitations could impair the second choice, or participants may set a different speed-accuracy trade-off between the first and second choices.

### 6. Task and position biases

#### 6.1. Task-related bias

Apart from confidence, other factors could have influenced choice order in our experiments. For instance, biases related to the nature of the task. As Figure 2A and Figure S3B show, some participants were overall more biased towards responding first to the color task, while some others were biased towards the letter task. For the three first experiments, a monotonic measure of bias magnitude could be obtained by averaging the proportion of trials where color was chosen first across trial types, and taking the absolute value of this average. This bias magnitude could be related to one of the measures of the priority effect. For Figure S6 we chose to relate it to the difference of proportions of trials where the color task was chosen first between EH and HE trials (the same measure shown for experiments 1, 2 and 3 in figures S3C, 2B and 2D, respectively). A Pearson correlation between the color/letter bias magnitude and the priority effect yielded a significant negative relationship (*r* = −0.370, *p* = 0.001), which suggests that a bigger task bias decreases the priority effect based on confidence.

**Figure S6.**
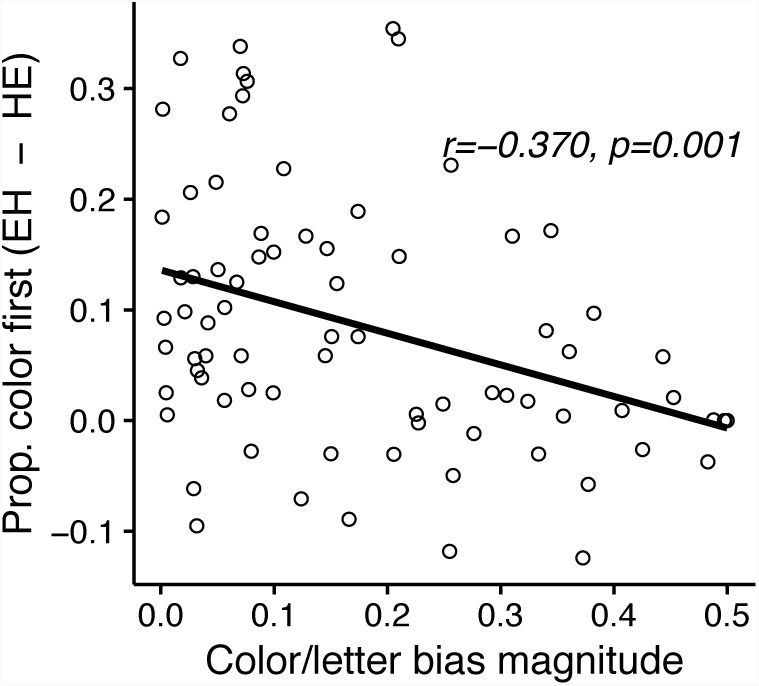
For experiments 1, 2 and 3, difference between proportion color task chosen first for EH trials and proportion color task chosen first for HE trials, as a function of the magnitude of the color/letter bias. Each dot represents a participant. The inset text shows the *r* and *p* of a Pearson correlation between the variables on each axis, and the solid, black line on the plot represents the respective linear fit of the data.

#### 6.2. Position-related bias

In our experiments, another possible bias has to do with the position of response boxes. Some participants may have favored certain positions on the screen when making their choices by clicking on the response box occupying that position. Figure S7 plots, for all 5 experiments, the proportion of trials for which the response occupying each of the possible screen positions was chosen first. For the first three experiments, a two-way repeated measures ANOVA was run, with the proportion of trials chosen as a dependent variable, and with vertical (top-bottom) and horizontal (left-right) position of the response boxes as factors. For Experiment 1 the main effect of vertical position was significant (*F*(1,27) = 24.387, *p* < 0.001, 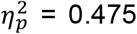), with horizontal position (*p* = 0.371) and interaction between both (*p* = 0.239) being non-significant. For Experiment 2 vertical position reached significance (*F*(1,19) = 11.460, *p* = 0.03, 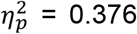), with horizontal position (*p* = 0.309) and interaction (*p* = 0.105) being non-significant. For Experiment 3, both the main effects of vertical (*F*(1,29) = 30.020, *p* < 0.001, 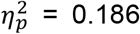) and horizontal (*F*(1,29) = 6.630, *p* = 0.015, 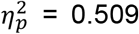) position were significant, but not their interaction (*p* = 0.997). Overall, there was a tendency to favor top response boxes when making the first choice. In the case of Experiment 4, we could explore a possible position bias when choosing which set of trials to face first. That bias would either favor the left or the right choice boxes. The proportion of blocks for which the left box was chosen was significantly bigger than 0.5 (t(30) = 2.352, *p* = 0.025, 95% CI [0.512, 0.668], *d* = 0.422), thus revealing a tendency to choose the left box. The same analysis was repeated for Experiment 5, revealing a non-significant trend in the same direction (t(28) = 1.908, *p* = 0.067, 95% CI [0.495, 0.636], *d* = 0.354). A similar analysis could be done for Experiment 6, now with the proportion of problems for which the left box was chosen. This was significantly bigger than 0.5 (t(100) = 7.840, *p* < 0.001, 95% CI [0.635, 0.727], *d* = 0.855), again showing a bias towards choosing the left box.

**Figure S7.**
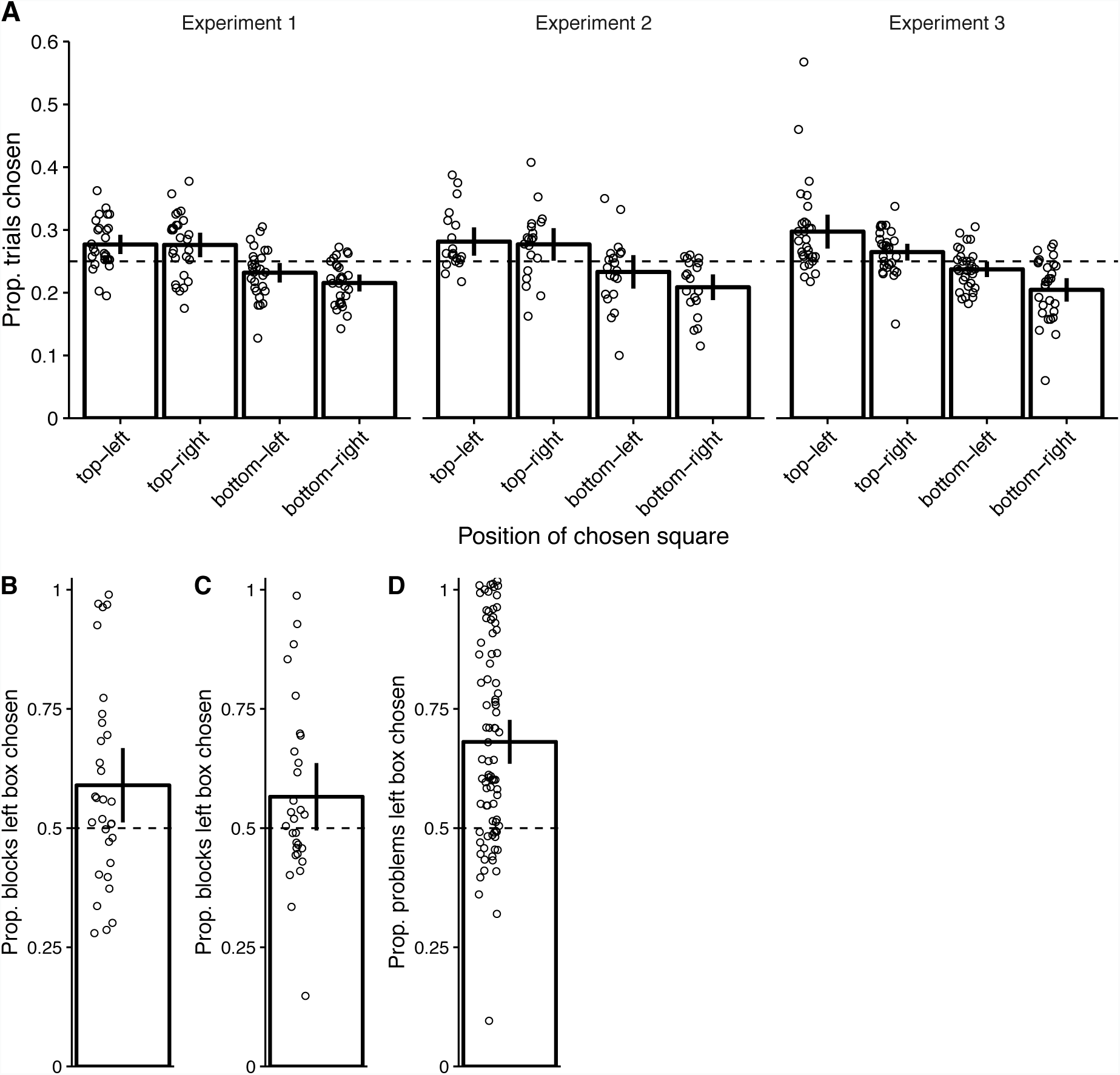
**A.** For experiments 1, 2 and 3, for each of the four possible positions response boxes could have, proportion of trials where the first choice was made on the choice box occupying that position. Bars represent across-participant averages, with error bars showing 95% confidence intervals. Each participant’s average is displayed with a single dot. The horizontal, dashed line marks a proportion of 0.25, corresponding to a non-biased ideal. **B.** For Experiment 4, proportion of blocks where the chosen set occupied the left choice box. The bar represents across-participant averages, with the error bar showing 95% confidence interval. Each participant’s average is displayed with a single dot. The horizontal, dashed line marks a proportion of 0.5, corresponding to a non-biased ideal. **C.** Same as B, but for Experiment 5. **D.** For Experiment 6, proportion of problems where the chosen problem occupied the left choice box. The bar represents across-participant averages, with the error bar showing 95% confidence interval. Each participant’s average is displayed with a single dot. The horizontal, dashed line marks a proportion of 0.5, corresponding to a non-biased ideal.

### 7. Effect of an easy-first approach on performance

#### 7.1. Experiments 1 to 5

After finding an easy-first effect in our data, a logical follow-up question is whether dealing first with easy tasks improves performance as compared to dealing first with difficult tasks. We first tackled this question by analyzing previous data. We took the dataset from experiments 1 to 3 and selected only those trial types where the two tasks had different difficulty (EH and HE). We then split that subset of data between those trials where the easy task had been chosen first and those where the hard task had been chosen first. Then we calculated the average observed performance per participant, but we did so split by the expected difficulty of each task within a trial, in order to appreciate possible differences between easy and hard tasks. This is illustrated in Figure S8A. We ran a two-way repeated measures ANOVA, where the dependent variable was observed performance and the factors were the expected difficulty of the task chosen first, and expected difficulty of the task at hand. As expected, the main effect of task difficulty was significant for all experiments (Experiment 1: *F*(1,27) = 1097.485, *p* < 0.001, 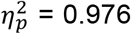; Experiment 2: *F*(1,19) = 481.122, *p* < 0.001, 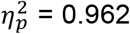; Experiment 3: *F*(1,29) = 912.556, *p* < 0.001, 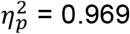). On the other hand, the main effect of difficulty of the task chosen first was not significant for the first two experiments (Experiment 1: *p* = 0.911; Experiment 2: *p* = 0.150), reaching marginal significance for Experiment 3 (*F*(1,29) = 3.579, *p* = 0.069, 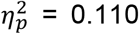). Interestingly, the interaction between the main two factors was significant for Experiment 1 *(F*(1,27) = 17.278, *p* < 0.001, 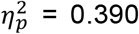) and Experiment 2 (*F*(1,19) = 7.394, *p* = 0.014, 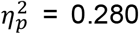), and marginally significant for Experiment 3 (*F*(1,29) = 3.527, *p* = 0.071, 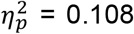). As Figure S8A helps to illustrate, the direction of this interaction seemed to imply that, when the task was hard, responding to it first improved performance, as compared to when it was responded to first. The same happened when the task was easy: performance on it was better when responded to first. Thus, independently of the difficulty, responding to a task first seemed to bring an advantage in performance on that particular task, but there was no overall advantage behind responding always easy-first or hard-first.

We conducted a similar analysis for Experiment 4. However, each trial had only one task, so the variables we worked with were different. We took all the trials from the test phase and, for each participant, we calculated average performance, grouping trials by their set’s expected difficulty (easy vs. hard) and by the expected difficulty of the set that the participant chose to do first on that block (easy vs. hard). This can be seen in Figure S8B We then used both variables as factors and performance as dependent variable for a two-way repeated measures ANOVA. The only significant main effect was expected difficulty (*F*(1,30) = 924.600, *p* < 0.001, 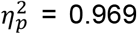), with the difficulty of the chosen set (*p* = 0.981) and the interaction between both main effects (*p* = 0.857) not reaching significance. Thus, we found no evidence of any impact on performance of choosing certain difficulty first.

In Experiment 5 both conditions had the same expected difficulty. Thus, we could not look into a possible advantage of tackling first the easy task. However, we could still investigate whether prioritizing the low condition led to a better performance. We ran a two-way repeated measures ANOVA with average observed performance for the test phase as dependent variable. As factors, we used that trial’s condition, and the condition that block’s prioritized set belonged to. No main effects reached significance (trial’s condition: *p* = 0.578; block’s prioritized condition: *p* = 0.912), and the interaction between both was not significant either (*p* = 0.494). Figure S8C helps see the absence of a relationship between any prioritization and performance.

We finally focused on Experiment 6. Given the ceiling effect on performance for this experiment, though, we used response time as the dependent variable. For each participant, we calculated median response times (in order to downweight outliers corresponding to trials where the participant took a considerable time to respond) per problem, grouping problems within a trial according to their expected difficulty, and to the expected difficulty of the problem the participant chose to do first on that block (see Figure S8D). However, in order to have data from every participant on each category, we had to remove 4 participants who always chose the easy problem first. After running an ANOVA, the main effect of the problem’s difficulty reached significance (*F*(1,79) = 24.570, *p* < 0.001, 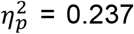), but not the main effect of the difficulty of the chosen problem (*p* = 0.189), or their interaction (*p* = 0.516). Response times, then, were not affected by an easy-first approach.

**Figure S8.**
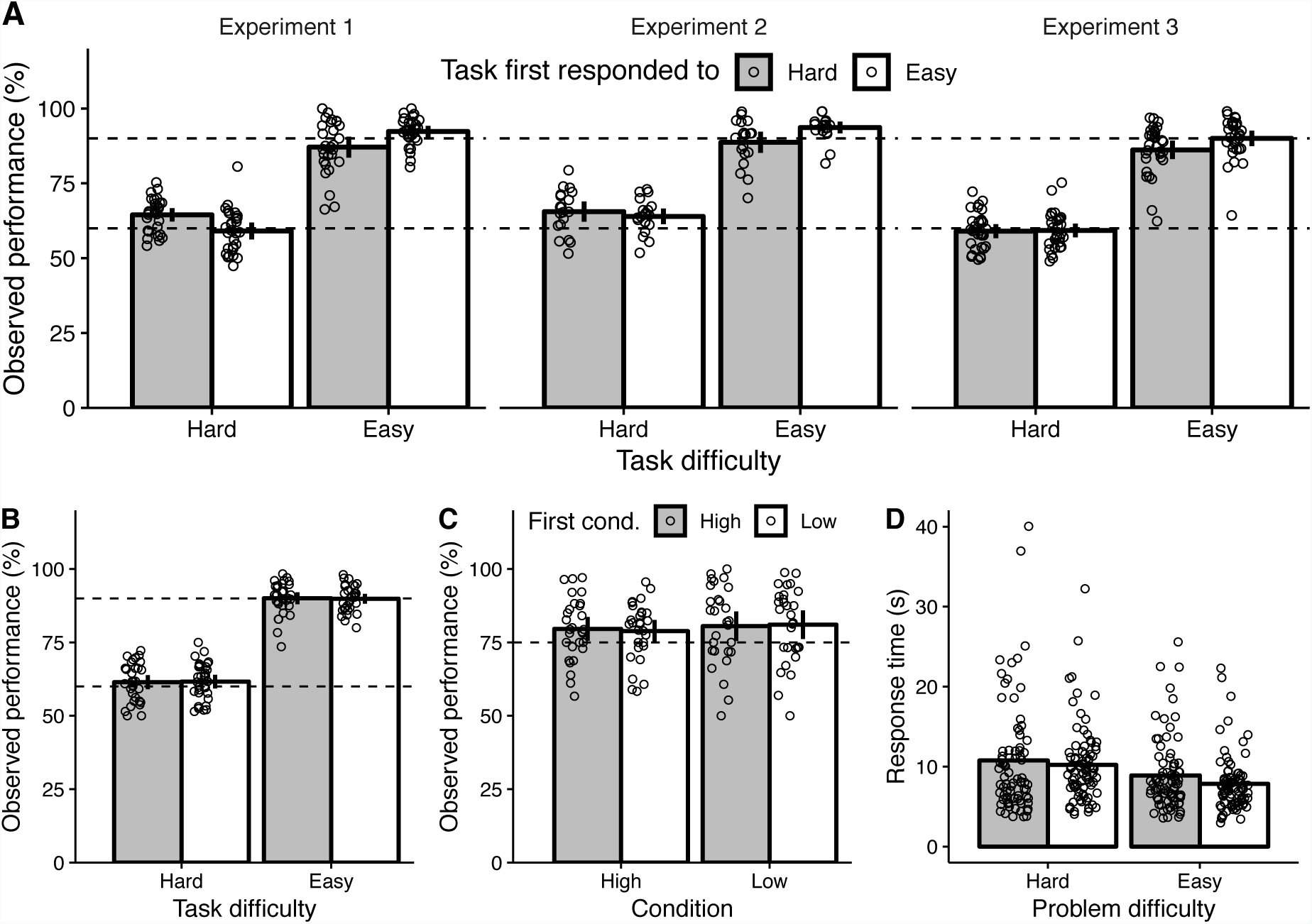
**A.** For experiments 1 to 3 (different panels), observed performance (as a percentage) as a function of the expected difficulty of the task (x-axis) and the task first responded to (color-coded). Bars represent averages across participants, with error bars denoting 95% confidence intervals. Dots represent individual participants. Only data from the EH and HE trial types was used. Top and bottom dashed lines help indicate where 90% and 60% observed performance would lay, respectively. **B.** Same as A, but for Experiment 4. Only trials from the test phase are used. However, the task first responded to refers to the whole block. **C.** For Experiment 5, observed performance (as a percentage) as a function of the condition for that trial (x-axis) and the condition whose set was prioritized in that block (color-coded). Bars represent averages across participants, with error bars denoting 95% confidence intervals. Dots represent individual participants. The dashed line indicates where 75% observed performance would lay.Only trials from the test phase are used. **D.** For Experiment 6, response time as a function of expected difficulty of the problem (x-axis) and the problem first responded to (color-coded to match the rest of the figure). Dots represent the median response times of individual participants, and bars represent the average of those medians.

#### 7.2. Supplementary experiment controlling response order

We decided to run an experiment with a new sample (n=16), where the central manipulation consisted in controlling response order. In half the trials response order was free, exactly as in experiments 1 to 3. In half the other trials, though, response order was imposed. After the stimulus, only two choice boxes appeared on the screen: either those related to the color task, or those related to the letter task. After a choice was made, the two boxes were replaced by those two corresponding to the other task, and participants had to make the second choice. Half the trials with imposed order required responding to color and then to letter, and vice versa in the other half. Within a trial, the difficulty of the two tasks was always different: thus, trials were only either EH or HE, with half the trials within a block being of each trial type. Participants completed 4 parts of 4 blocks each, with each block having 24 trials. That number was chosen so, within a block, the same trial type (EH or HE) and response order (imposed easy first, imposed hard first, or free, with this one being presented as many times as both imposed orders together) were presented at the same ratio, although in a random order. As for the payoff system, it was exactly as in Experiment 2.

Performance in this experiment is illustrated in Figure S9A. We ran a three-way ANOVA with observed performance as dependent variable, and where the three factors were response order condition (free vs. imposed), task difficulty (easy vs. hard) and task responded to first (easy vs. hard). As expected, the main factor of task difficulty was highly significant (*F*(1,15) = 616.9, *p* < 0.001, 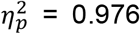), but none of the other factors nor interactions reached statistical significance (condition: *p* = 0.929; task responded to first: *p* = 0.923; condition x task difficulty: *p* = 0.788; condition x task responded to first: *p* = 0.669; task difficulty x task responded to first: *p* = 0.414; triple interaction: *p* = 0.188).

In order to further strengthen the main claim of our study, we also used the data from the present experiment to find support for priority based on confidence. As already done with previous experiments, we chose to use expected difficulty as a proxy for confidence. When comparing the two trial types of this experiment, we found that the color task was responded to first on 54% of EH trials and on 49% of HE trials. The difference of proportions, plotted in Figure S9B, was significant across participants (t(15) = 2.467, *p* = 0.026, 95% CI [0.007, 0.100], *d* = 0.617), thus providing further support for our central message.

**Figure S9.**
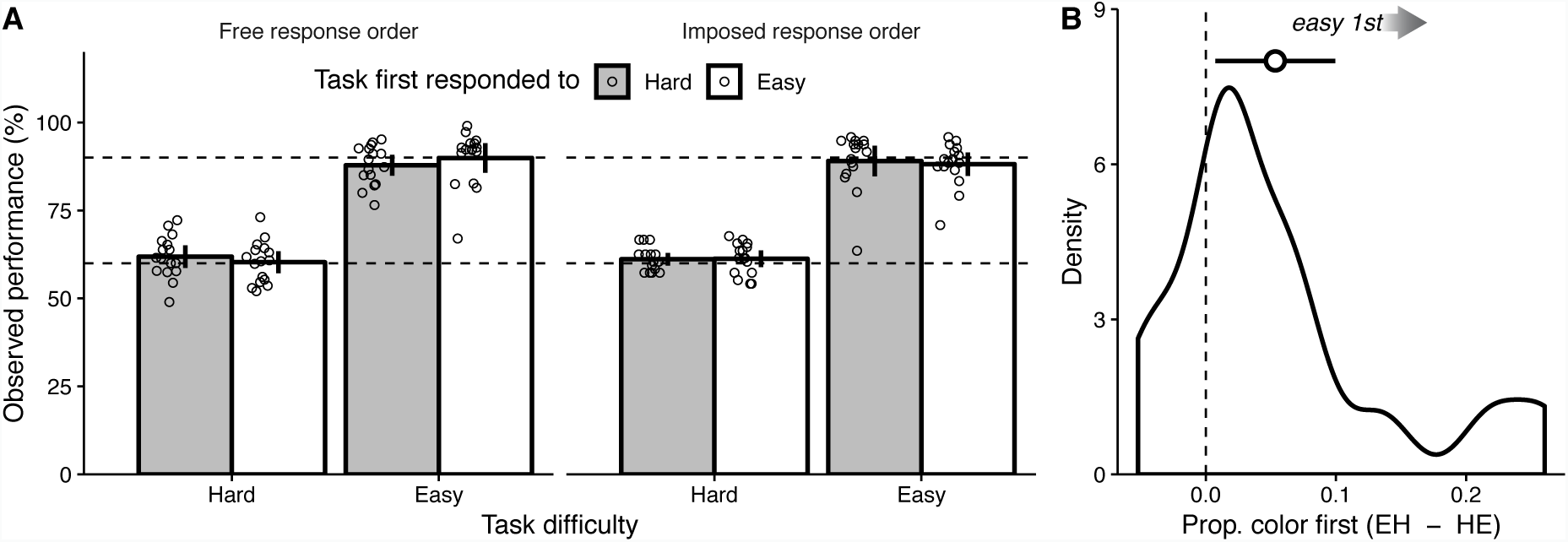
Supplementary experiment controlling response order: results. **A.** For both response order types (different panels), observed performance (as a percentage) as a function of expected difficulty of the task (x-axis) and the task first responded to (color-coded). Bars represent averages across participants, with error bars denoting 95% confidence intervals. Dots represent individual participants. Top and bottom dashed lines help indicate where 90% and 60% observed performance would lay, respectively. **B.** Density of the difference between proportion color task chosen first for EH trials and proportion color task chosen first for HE trials. The density is drawn by using the difference of proportions for each participant. The across-participant average is displayed as a big dot with error bars showing 95% confidence intervals. The dashed line corresponds to no difference between EH and HE trials. The sample size for this experiment was 16 participants, each participant completing 384 trials.

### 8. Supplementary performance analyses for Experiment 5

The present section offers a more in-depth analysis of performance in Experiment 5. We compare performance among different parts or phases of our experiment and offer figures to illustrate some of the information given in the main body of the manuscript.

In the main text we showed that performance was not different between the low and the high conditions (see also Figure 4B). This analysis was conducted over all trials across different parts of the experiment: the familiarization and test phases of the main part, and the final trials where confidence had to be reported. We confirmed that performance was also similar between the low and high conditions when analyzing separately the familiarization phase (t(28) = −1.027, *p* = 0.313, 95% CI [-0.087, 0.029], *d* = −0.191), the test phase (t(28) = −0.468, *p* = 0.643, 95% CI [-0.070, 0.044], *d* = −0.087), and the confidence trials (t(28) = −1.281, *p* = 0.211, 95% CI [-0.100, 0.023], *d* = −0.238). This is illustrated in Figure S10A.

Also in the main text we showed the relationship between the greater confidence in the low condition and the greater tendency to prioritize low condition sets. This relationship could not be explained by a performance advantage in low condition trials, as previously reported, as no significant correlation was found between the difference of performance between low and high trials and the proportion of blocks where the low condition set was chosen first (*r* = - 0.099, *p* = 0.611). This is illustrated in Figure S10B.

Finally, we made sure that performances among the different parts of the experiment were similar within participants. This is important, since in our study we link performance in the main part of the experiment with confidence in the final trials where it had to be reported. We found that across participants, performance in the familiarization trials was correlated with performance in the test trials, both for the high condition (*r* = 0.820, *p* < 0.001) and for the low condition (*r* = 0.928, *p* < 0.001). Figure S10C illustrates this. We then pooled all the trials of the main part together and ran a correlation with performance on the final trials where confidence had to be rated (see Figure S10D). The correlation was significantly positive for both conditions (high: *r* = 0.720, *p* < 0.001; low: *r* = 0.831, *p* < 0.001).

**Figure S10.**
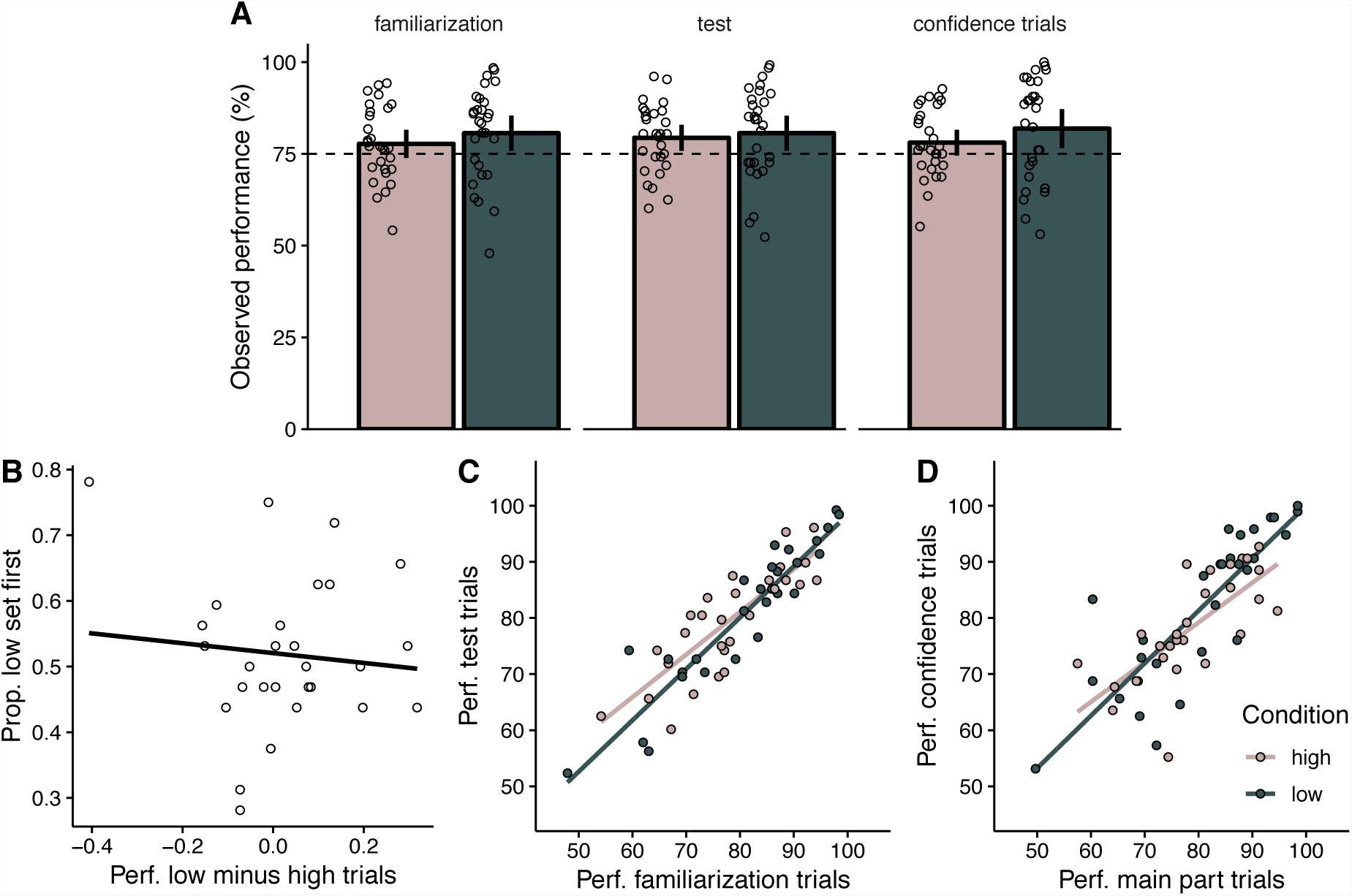
Supplementary performance analyses for Experiment 5. **A.** For the familiarization and test phases of the main part, and for the final trials where confidence reports were required, observed performance as a function of expected performance, both in percentage, split by condition. Bars represent averages across participants, with error bars denoting 95% confidence intervals. Dots represent individual participants. The horizontal dashed line indicates where 75% observed performance would lay. **B.** Proportion of blocks where participants chose to complete first the the set whose trials belonged to the low condition, as a function of the difference, for the familiarization phase, between the average performance for low condition trials minus the average performance for high condition trials. The line depicts a linear fit of the data. **C.** Average performance for the familiarization trials against average performance for the test trials. Each participant is depicted with two dots, one corresponding to each condition (color-coded). Lines represent linear fits for each condition’s data. **D.** Average performance for the main part of the experiment (with the familiarization and test trials being pooled together) against average performance for those trials where confidence had to be rated. Each participant is depicted with two dots, one corresponding to each condition (color-coded). Lines represent linear fits for each condition’s data.

### 9. Stimulus materials

#### 9.1. Complete list of animal names used for experiments 4 and 5

The following list contains all the animal names used to label sets in experiments 4 and 5

. The original French name is written first in capital letters, as displayed in the experiment, followed by the English translation (between brackets).

|  |  |  |
| --- | --- | --- |
| ABEILLE (bee) | AGNEAU (lamb) | AIGLE (eagle) |
| ANTILOPE (antelope) | AUTRUCHE (ostrich) | BABOUIN (baboon) |
| BALEINE (whale) | BISON (bison) | BUFFLE (buffalo) |
| CANARD (duck) | CASTOR (beaver) | CHAMEAU (camel) |
| CHENILLE (caterpillar) | CHEVAL (horse) | CHÈVRE (goat) |
| CHIEN (dog) | CHOUETTE (barn owl) | CIGOGNE (stork) |
| COCHON (pig) | COYOTE (coyote) | CRIQUET (cricket) |
| DAUPHIN (dolphin) | ÉCUREUIL (squirrel) | ÉLÉPHANT (elephant) |
| ESCARGOT (snail) | FOURMI (ant) | FURET (ferret) |
| GAZELLE (gazelle) | GIRAFE (giraffe) | GORILLE (gorilla) |
| GUÉPARD (cheetah) | HAMSTER (hamster) | HERISSON (hedgehog) |
| HIBOU (owl) | HYENE (hyena) | JAGUAR (jaguar) |
| LAPIN (rabbit) | LIMACE (slug) | LOUTRE (otter) |
| LUCIOLE (firefly) | MARMOTTE (groundhog) | MOINEAU (sparrow) |
| MOUTON (ram) | OTARIE (seal) | PANDA (panda) |
| PAPILLON (butterfly) | PIGEON (pigeon) | POISSON (fish) |
| PONEY (pony) | POULE (chicken) | RENARD (fox) |
| RENNE (reindeer) | REQUIN (shark) | SANGLIER (boar) |
| SCARABÉE (beetle) | SERPENT (snake) | SINGE (ape) |
| SOURIS (mouse) | TAUREAU (bull) | TIGRE (tiger) |
| TORTUE (turtle) | VACHE (cow) | VAUTOUR (vulture) |
| ZEBRE (zebra) |  |  |

#### 9.2. Complete list of arithmetic problems used for Experiment 6

The following list shows the schematics, actual problem and answer for each pair of easy and hard arithmetic problems used in Experiment 6. The two problems within a pair (each row) were always presented together.

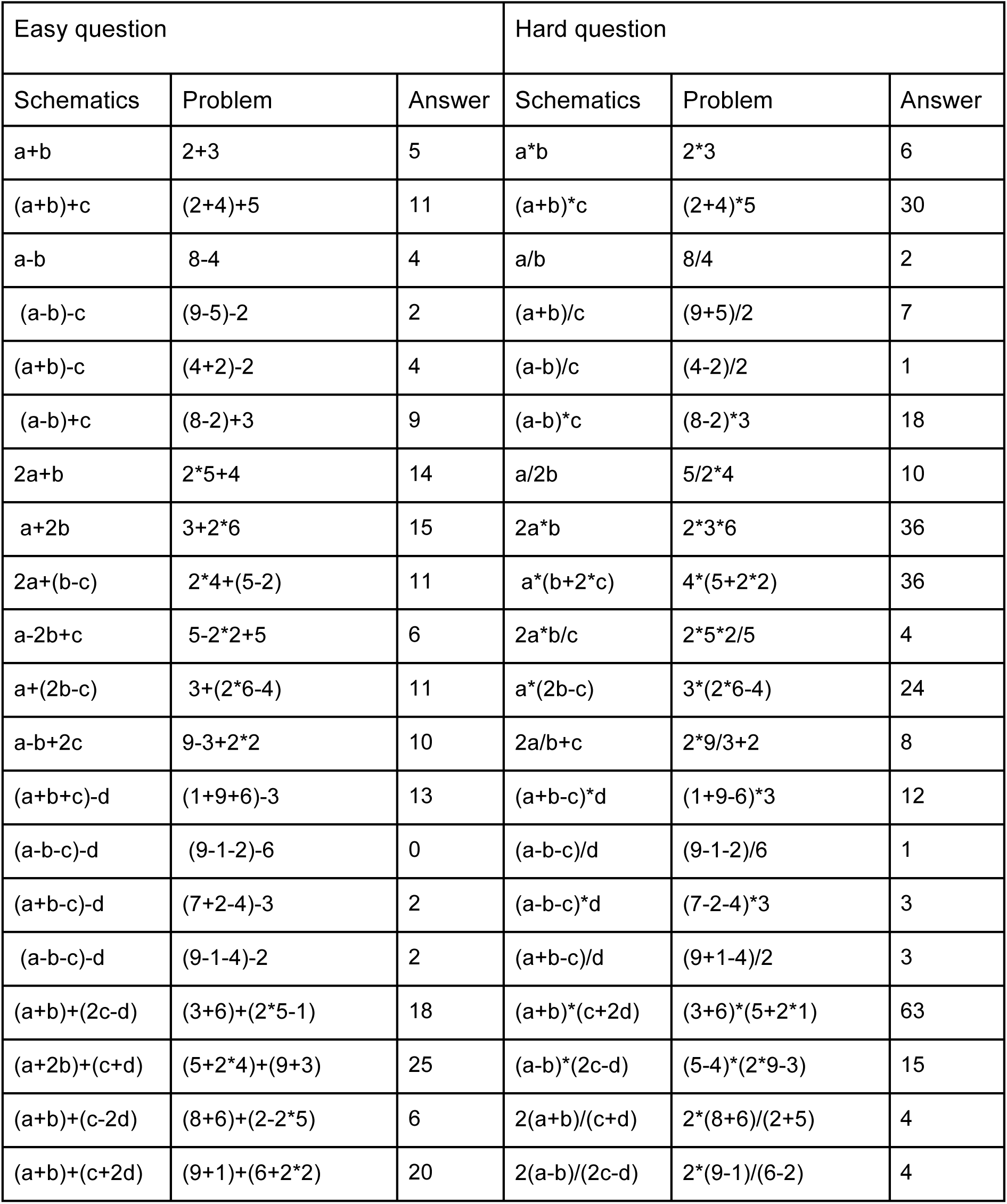

| Easy question |  |  | Hard question |  |  |
| --- | --- | --- | --- | --- | --- |
| Schematics | Problem | Answer | Schematics | Problem | Answer |
| $a+b$ | $2+3$ | 5 | $a*b$ | $2*3$ | 6 |
| $(a+b)+c$ | $(2+4)+5$ | 11 | $(a+b)*c$ | $(2+4)*5$ | 30 |
| $a-b$ | $8-4$ | 4 | $a/b$ | $8/4$ | 2 |
| $(a-b)-c$ | $(9-5)-2$ | 2 | $(a+b)/c$ | $(9+5)/2$ | 7 |
| $(a+b)-c$ | $(4+2)-2$ | 4 | $(a-b)/c$ | $(4-2)/2$ | 1 |
| $(a-b)+c$ | $(8-2)+3$ | 9 | $(a-b)*c$ | $(8-2)*3$ | 18 |
| $2a+b$ | $2*5+4$ | 14 | $a/2b$ | $5/2*4$ | 10 |
| $a+2b$ | $3+2*6$ | 15 | $2a*b$ | $2*3*6$ | 36 |
| $2a+(b-c)$ | $2*4+(5-2)$ | 11 | $a*(b+2*c)$ | $4*(5+2*2)$ | 36 |
| $a-2b+c$ | $5-2*2+5$ | 6 | $2a*b/c$ | $2*5*2/5$ | 4 |
| $a+(2b-c)$ | $3+(2*6-4)$ | 11 | $a*(2b-c)$ | $3*(2*6-4)$ | 24 |
| $a-b+2c$ | $9-3+2*2$ | 10 | $2a/b+c$ | $2*9/3+2$ | 8 |
| $(a+b+c)-d$ | $(1+9+6)-3$ | 13 | $(a+b-c)*d$ | $(1+9-6)*3$ | 12 |
| $(a-b-c)-d$ | $(9-1-2)-6$ | 0 | $(a-b-c)/d$ | $(9-1-2)/6$ | 1 |
| $(a+b-c)-d$ | $(7+2-4)-3$ | 2 | $(a-b-c)*d$ | $(7-2-4)*3$ | 3 |
| $(a-b-c)-d$ | $(9-1-4)-2$ | 2 | $(a+b-c)/d$ | $(9+1-4)/2$ | 3 |
| $(a+b)+(2c-d)$ | $(3+6)+(2*5-1)$ | 18 | $(a+b)*(c+2d)$ | $(3+6)*(5+2*1)$ | 63 |
| $(a+2b)+(c+d)$ | $(5+2*4)+(9+3)$ | 25 | $(a-b)*(2c-d)$ | $(5-4)*(2*9-3)$ | 15 |
| $(a+b)+(c-2d)$ | $(8+6)+(2-2*5)$ | 6 | $2(a+b)/(c+d)$ | $2*(8+6)/(2+5)$ | 4 |
| $(a+b)+(c+2d)$ | $(9+1)+(6+2*2)$ | 20 | $2(a-b)/(2c-d)$ | $2*(9-1)/(6-2)$ | 4 |

